# Subunit Vaccination Using Atomic Layering Thermostable Antigen and Adjuvant (ALTA^®^) Platform Elicits Enhanced Humoral and Cellular Immune Responses

**DOI:** 10.64898/2026.01.05.697739

**Authors:** Daria L. Ivanova, Matthew S. Lewis, Annie B. Caplan, Isabella R. Walters, Emma M. Snyder, Keith A. Strand, Lorena R. Antunez, Sineenart Sengyee, Sarah B. Weiby, Federico Urbano-Munoz, Mary N. Burtnick, Paul J. Brett, Sky W. Brubaker

## Abstract

Creating effective and thermostable vaccines is of significant relevance for public health. The Atomic Layering Thermostable Antigen and Adjuvant (ALTA^®^) platform combines spray drying to stabilize antigens in a sugar matrix followed by coating with atomic layer deposition (ALD) for temporal control over *in vivo* release. While these technologies have shown preliminary promise for different vaccine antigens, further characterizations of the immune response to ALTA^®^ formulated antigens are still needed. Here, the immune response to ALTA^®^ formulated antigens is described and compared to a set of adjuvanted liquid vaccine formulations that included Alhydrogel^®^, AddaVax^™^, and Alhydrogel^®^+CpG. The humoral and cell-mediated responses were measured by ELISA and flow cytometry. Increased and lasting antigen-specific antibody titers following administration of ALTA^®^ containing ovalbumin (OVA) demonstrated robust and durable humoral response. Furthermore, ALTA^®^ injected mice produced both IgG2c and IgG1 indicating a balanced Th1/Th2 response. Importantly, ALTA^®^ OVA elicited robust humoral response at lower doses of aluminum than Alhydrogel^®^. The most striking difference between ALTA^®^ and the liquid vaccine formulations tested was a greater OVA-specific CD8+ T cell response observed at all antigen doses tested. Mechanistically, antigen encapsulation within ALTA^®^ particles was critical for antibody production and CD8+ T cell responses as well as antigen capture by antigen-presenting cells (APCs) at the site of injection and draining lymph nodes. To test these concepts in a more physiological application, protein and polysaccharide vaccine antigens derived from a facultative intracellular bacterium *Burkholderia pseudomallei*, the causative agent of melioidosis, were formulated using the ALTA^®^ platform. Compared to liquid vaccine formulations, ALTA^®^ immunized mice showed enhanced antigen-specific antibody production and IFN-γ secreting T cell responses using lower adjuvant doses of aluminum and CpG. Overall, ALTA^®^ formulated protein and polysaccharide antigens elicited strong humoral and cell-mediated immunity suggesting potential broad applicability of the platform to vaccines against various diseases, including against cancer and infections from intracellular pathogens.

## 1. Introduction

Currently several licensed vaccines require continuous and robust cold-chain supply management that limits their distribution and availability across the globe. In addition, vaccines generally require multiple doses to elicit lasting immunity. Often patient compliance is low for booster doses resulting in poor vaccine coverage. Therefore, thermostable vaccine technologies enabling improved efficacy using fewer administrations are of significant relevance for global health.

The Atomic Layering Thermostable Antigen and Adjuvant (ALTA^®^) platform technology is an approach that combines spray drying to incorporate antigens in a sugar matrix followed by atomic layer deposition (ALD) generating an alumina coating for temporal control over *in vivo* release^1^. Embedding of vaccine components within glassy organic matrices formed from disaccharide containing mixtures via lyophilization or spray drying can protect antigens and adjuvants from physical and chemical degradation^2,3^. Importantly, lyophilized or spray dried antigens and adjuvants maintain their immunogenicity^4^. For example, spray dried formulations of human papillomavirus (HPV) type 16 L1 capsomeres with glass-forming polymers and trehalose maintained the capacity to induce antigen-specific antibody production in mice^4^. The second technology utilized in the ALTA^®^ platform is ALD^5^. During this process, spray dried particles are treated with iterative cycles of ALD adding surface alumina (Al_2_O_3_) to achieve different coat thicknesses. Increasing the coat thickness delays the timing of antigen release *in vitro* and antibody production *in vivo*^1^. Thus, the number of ALD cycles can be adjusted and combined with a priming dose to create a single administration product with the potential to replace multiple administrations of multi-dose vaccines^6,7^. Recently, a single administration of mosaic-8b receptor-binding domain nanoparticles prepared using ALD technology elicited antibodies with improved mismatched binding and neutralization of SARS-like betacoronaviruses (sarbecoviruses), compared to conventional prime-boost immunizations^7^. Additionally, a single administration of ALD-coated Rabies virus (RABV) vaccine induced IgG against Rabies G protein and virus neutralizing antibody titers significantly higher than those generated in response to three administrations of conventional liquid RABV vaccine^6^.

Previous research has shown that spray dried and ALD-coated vaccine formulations can increase the production of binding and neutralizing antibodies over formulations without alumina coating^1,4,7,8^. The mechanisms contributing to improved immunogenicity for ALD-coated vaccine formulations are not fully understood but may include the following. First, alumina layers may protect the spray dried vaccine antigens/adjuvants in its core during storage^9^. Second, alumina layers may protect antigens from rapid *in vivo* processing thereby extending bioavailability of intact antigens. Prolonging the presence of native antigens *in vivo* improves the germinal center (GC) response in the draining lymph nodes (dLN) and promotes improved humoral immunity^10–13^. Third, the alumina layers may have an intrinsic adjuvating effect, similar to other aluminum-containing adjuvants, such as Alhydrogel^®^ and Adju-Phos^®14,15^. Moreover, physical characteristics of the alumina-coated microparticles, including their spherical shape, surface charge, and size, may promote their uptake by APCs^8^. A general but defining feature of an effective vaccine is the ability to drive antigen trafficking from the injection site to the dLN^16^. Thus, ALD-coated vaccines may promote antigen delivery to dLN via uptake by APCs. Clearly further investigation is needed given the number of hypotheses regarding the mechanism(s) contributing to improved ALD-coated vaccine immunogenicity.

To date, the immune response to ALD-coated vaccines, including ALTA^®^, has primarily been evaluated for humoral immunity via ELISA and neutralization assays^1,4,6–8^. Although vaccine efficacy is typically measured by an antibody-mediated response, a growing number of studies show that vaccine-elicited cellular immunity provides protection against severe disease and reduces mortality rates^17,18^. For example, T cell mediated immunity is critical for protection against rapidly mutating intracellular (HIV-1, influenza, coronaviruses, etc.), latent pathogens (HIV-1, VZV, etc.) and cancer ^19,20^. Since most effective vaccines are likely to employ both humoral and cellular immune responses, measuring the cell-mediated response to ALTA^®^ vaccines is important.

To properly evaluate the preclinical merit of a novel platform technology it can be useful to compare against the technologies that have been approved for use clinically. For vaccines, that primarily means that a comparison with an aluminum-based formulations is warranted. After the approval of aluminum hydroxide formulation as a vaccine adjuvant for human use in 1939, aluminum-based adjuvants have been widely used in many commercial vaccines, including diphtheria-tetanus-pertussis, human papillomavirus, and hepatitis^15^. Effective in inducing a strong Th2 response, classic aluminum salt adjuvants demonstrate a limited activation of the Th1-biased cell-mediated immunity, which restricts their use in vaccines against intracellular pathogens, including tuberculosis, malaria, and HIV-1^21^. Other adjuvants have shown greater effectiveness in eliciting cell-mediated immune responses. An emulsion-based squalene oil-in-water based adjuvant, MF59, became the first non-aluminum salt adjuvant to be approved for human use^22^. Compared to aluminum-salt adjuvants, MF59 demonstrated a more efficient recruitment of APCs, trafficking of antigen-loaded APCs to dLNs, and activation of both cellular and humoral responses^23,24^. In addition, toll-like receptor (TLR) agonists are commonly used to induce a strong Th1-biased cellular immune response characterized by increased IFN-γ production and activation of cytotoxic T lymphocytes. To reduce adverse reactions, TLR agonists are typically adsorbed to aluminum salts. A synthetic analog of bacterial DNA CpG1018 TLR9 agonist is used in an approved HBV vaccine. Collectively, the adjuvants used in clinically approved vaccine formulations can provide a benchmark by which to measure the immunogenicity of the ALTA^®^ platform against.

The number of disease-specific vaccine antigens formulated using spray drying and ALD technologies is rapidly growing^4,6–8,25–28^. Here, the ALTA^®^ technology was applied to antigens derived from a facultative intracellular Gram-negative bacterium *Burkholderia pseudomallei*, the causative agent of melioidosis^29^. Despite challenges in diagnosis and treatments resulting in high morbidity and mortality rates, there are currently no licensed vaccines against this disease^29^. Further complicating vaccine development efforts is the fact that *Burkholderia* is historically endemic to tropical regions of Southeast Asia and Northern Australia, where the requirements for cold-chain vaccine storage are more challenging. A lead subunit vaccine candidate has shown promise by providing protection in mice^30,31^. This candidate vaccine contains highly conserved antigens expressed by *B. pseudomallei* and *B. mallei*, specifically the 6-deoxyheptan capsular polysaccharide (CPS) conjugated to the carrier protein CRM197 and the T6SS-1 associated hemolysin coregulated protein 1 (Hcp1). While vaccine protection correlates with high anti-CPS IgG titers and Hcp1-specific IFN-γ T cell responses^30^, protection against inhalational challenge required multiple (2-3) doses of this subunit vaccine adjuvanted with Alhydrogel^®^ and CpG (ODN 2006 (ODN 7909))^29^. Therefore, there is tremendous interest in the development of a thermostable, single-dose vaccine against *B. pseudomallei*.

In this paper, immune responses were characterized for vaccines formulated using the ALTA^®^ platform and compared against liquid vaccine formulations containing commonly used adjuvants known to elicit strong humoral and/or cellular immunity. The study demonstrates immunogenicity by measuring antibody titers and T cell responses following administrations of OVA antigen formulated with the ALTA^®^ platform or with 1) Alhydrogel^®^, 2) AddaVax^™^ (mimetic of MF59), or 3) combined Alhydrogel^®^ and ODN 1018 (mimetic of Dynavax’s CpG 1018). Critical product attributes as well as the biological mechanisms by which immunogenicity is imparted by the ALTA^®^ platform were also explored. Finally, the *B. pseudomallei* vaccine candidates, CPS-CRM197 and Hcp1 were formulated with CpG ODN 2006 using ALTA^®^ platform and compared against liquid controls. In total, the studies were designed to further characterize or define the humoral and cellular immune responses elicited by the ALTA^®^ vaccine platform.

## 2. Materials and Methods

### 2.1. Manufacture of ALTA^®^ materials

Formulations were generated using a proprietary composition. Briefly, the formulations containing the model antigen ovalbumin (OVA) (InvivoGen, San Diego, CA, USA, #vac-pova-100) were made to contain 1% w/w OVA in the spray dried powder. The *B. pseudomallei* antigens CPS-CRM197 and Hcp1 were manufactured and provided by Mary Burtnick and Paul Brett of the University of Nevada, Reno. These antigens were formulated to create spray dried powders which contained 0.5-0.76 %w/w Hcp1, 0.45-0.69 %w/w CPS-CRM197, and 0.25-0.38 % w/w CpG 2006 (Genscript Piscataway, NJ). Intermediate spray dried powders were made using a Buchi Mini B-290 spray dryer with a B-296 Dehumidifier (BUCHI, New Castle, DE, USA). Spray drying setpoints were chosen to target the desired powder properties, including particle size, residual moisture content, and yield. ALD-coated powders were generated using the intermediate spray dried powders in a custom, mechanically agitated fluidized-bed ALD reactor, using trimethylaluminum and water as precursors^4,27^. Purge steps were employed after each precursor exposure to avoid chemical vapor deposition. The coating was performed at 50°C, using nitrogen as the fluidization gas. The number of ALD cycles used to coat powders in this study was 50.

The *Burkholderia* subunit vaccine candidates evaluated in the study included a CPS-CRM197 glycoconjugate combined with Hcp1 protein^30^. To produce CPS-CRM197, the 6-deoxyheptan CPS was purified from *B. thailandensis* BT2683 (an OPS-deficient derivative of strain E555), chemically activated, and covalently linked to recombinant, preclinical-grade recombinant CRM197 diphtheria toxin mutant (Fina Biosolutions) essentially as previously described^32^. The resulting CPS-CRM197 glycoconjugates used in this study contained 55% (wt/wt) CPS. Recombinant *B. mallei* hemolysin Hcp1 lacking a His-tag was purified from *E. coli* ^33^. Adjuvants included CpG (ODN 2006) oligonucleotide, which was synthesized by GenScript, and Alhydrogel 2% which was obtained from Invivogen.

To generate intentionally broken ALTA^®^ particles, powders were physically disrupted to break the alumina shell resulting in 86% immediately soluble fraction. The OVA concentration in broken ALTA^®^ was measured by SEC (determined to be 89 ug/mL, 91% of expected).

### 2.2. Animal Studies and Vaccine Administrations

All animal studies were conducted in accordance with the recommendations in the Guide for the Care and Use of Laboratory Animals. Animal studies were conducted at the University of Colorado, Boulder, or the University of Nevada, Reno. Study protocols were approved by UCB Institutional Animal Care and Use Committee (#2835, 2838) or UNR Animal Care and Use Committee (protocol no. 23-08-1194) and the U.S. Army Medical Research and Development Command Animal Care and Use Review Office (protocol no. CB11139.e001). Mice were housed in microisolator cages under pathogen-free conditions, provided with rodent feed and water ad libitum, and maintained on a 12-h light cycle.

For the ovalbumin antigen immunogenicity studies, female C57BL/6 mice aged 6–8 weeks were purchased from the Charles River Laboratories, Strain #000664. Vaccine administrations were given by intramuscular (i.m.) injection into the flank in a final volume of 50 μL. The dosing of ALTA^®^ OVA powder was based on the OVA antigen mass and delivered with saline containing 6% hydroxyethyl starch (Hospira NDC: 00409-7248-03). For immunizations with liquid OVA EndoFit™ (vac-pova, InvivoGen), the following compounds were used: Alhydrogel^®^ adjuvant 2% (1:50 OVA:Al3+, Vac-alu-50), AddaVax™ (vac-adx-10, InviVoGen), ODN 1018 VacciGrade™ (20 µg/dose, vac-1018-1), Poly(I:C) (HMW) VacciGrade™ (40 µg/dose, vac-pic, InVivoGen), αCD40 antibody (40 μg/dose; clone FGK4.5, BioXcell). The ODN 2006 (ODN 7909) was synthesized by GenScript. For immunizations with fluorescent OVA, the IVISense™ 680 NHS fluorescent dye (Revvity) was conjugated to OVA in house. In short, the OVA conjugation was done by reacting IVISense 680 NHS dye at a 10 molar ratio in 50 mM sodium borate pH 8 overnight. The conjugation was purified using desalting columns and dialysis to remove excess free dye. The degree of labeling (DOL) of this conjugation was 2.7 dye:OVA molecules. For ELISA, the serum was isolated from submandibular blood samples using gel separation tubes (Sarstedt, Nümbrecht, Germany, #41.1378.005) and stored at −80°C prior to analysis. For flow cytometry analysis, whole blood, spleen, dLN, or muscle tissue (injection site) were collected and processed immediately.

For the *Burkholderia* antigen immunogenicity studies, 6- to 8-week-old female C57BL/6 mice (Charles River Laboratories) were immunized subcutaneously (s.c.) with 50-coat ALTA^®^ particles or with liquid controls containing CPS-CRM197, Hcp1, Alhydrogel_®_, and CpG (doses detailed in Supplementary Figure 7). Liquid controls were formulated in PBS (pH 7.2) (Gibco). For the first study, mice (n = 5/group) were immunized with one dose of ALTA^®^ particles or liquid controls (Supplementary Figure 7A). After four weeks (day 28), serum and spleens were collected from terminally bled mice for use in ELISAs and ELISpot assays. For the second study, mice (n=10/group) were immunized with two doses (days 0 and 28) of ALTA^®^ particles or liquid controls containing high or matched adjuvant concentrations (Supplementary Figure 7B). Two weeks after the boost (day 42), serum and spleens were collected from terminally bled mice and used in ELISAs and ELISpot assays.

### 2.3. Quantitation of Antibody Titers by ELISA

#### Anti-OVA IgG1 and IgG2c

Immunized mouse serum samples were analyzed by an indirect, enzyme-linked immunosorbent assay (ELISA) to determine anti-OVA IgG1 and IgG2c antibody titers. Nunc 96-well flat bottom high-binding plates (Thermo Scientific, Norristown, PA, USA, #442404) were coated with 5 μg/well of OVA (Fisher, Hampton, NH, USA, #BP2535-5) in phosphate-buffered saline (PBS), incubated at 4°C overnight, and then rinsed four times using a wash buffer (0.05% Tween 20 in PBS). The plates were then incubated for 1 h at room temperature (RT) with an assay-blocking buffer (3% BSA and 0.05% Tween 20 in PBS). A standard curve was established with a dilution series of calibrated primary control (IgG1: Chondrex, Woodinville, WA, USA, #7093, IgG2c: Chondrex, #7109). The calibrated primary control and mouse sera samples were incubated for 1 h at RT, followed by a wash step to remove unbound antibody. Horseradish peroxide-conjugated secondary antibody (Southern Biotech, Birmingham, AL, USA, #1071-05 1:4000 dilution and Southern Biotech, #1078-05 1:4000 dilution) was added (50 μL/well) and incubated 1 h at RT. Excess secondary antibody was washed off, and the plates were developed for 20 min using Ultra TMB (Thermo Fisher, #34028) and quenched using sulfuric acid (H_2_SO_4_). Absorbance was measured on a BioTek Synergy plate reader (Agilent, Santa Clara, CA, USA) at 450 nm and 650 nm five minutes after the addition of H_2_SO_4_. The standard curves and interpolated data for each plate were independently generated using Gen5 software (Agilent, Santa Clara, CA, USA).

#### Anti-CPS and anti-Hcp1

Serum from terminally bled mice was obtained using Vacutainer SST tubes (BD) per the manufacturer’s instructions and stored at -80°C until required for use. Antibody responses directed against the CPS and Hcp1 were assessed by ELISAs as previously described^30^. Briefly, 96-well Maxisorp plates (Nunc) were coated overnight at 4°C with purified CPS or Hcp1 (1 μg/mL) solubilized in carbonate buffer (pH 9.6). The plates were blocked at room temperature for 30 min with StartingBlock T20 (Tris-buffered saline (TBS)) blocking buffer (Thermo Scientific) and then incubated for 1 h at 37°C with mouse serum samples serially diluted in TBS plus 0.05% Tween 20 (TBST; pH 7.5) plus 10% StartingBlock T20. To facilitate detection, the plates were incubated for 1 h at 37°C with 1/2,000 dilutions of anti-mouse IgG HRP-conjugated antibodies (Southern Biotech). The plates were developed with TMB (3,3′,5,5′-tetramethylbenzidine) substrate (KPL) and read at 620 nm using a FLUOstar Omega microplate reader (BMG Labtech). The reciprocals of the highest dilutions exhibiting optical densities (ODs) that were 3× background levels were used to determine the endpoint titers for the individual mice.

### 2.4. Cell Suspension for Flow Cytometry

Whole blood was collected into the EDTA-treated tubes (EDTA K3E, Sarstedt). Following the RBC lysis (ACK Lysing Buffer, Thermo Scientific) for 5 min RT, the blood samples were quenched, washed with complete RPMI (RPMI1640 with L-glutamate supplemented with 10% fetal bovine serum, penicillin/streptomycin, sodium pyruvate, non-essential amino acids, beta-mercaptoethanol, 4-(2-hydroxyethyl)-1-piperazineethanesulfonic acid (HEPES)), and plated onto 96-well U-bottom plates for staining.

For staining of splenocytes, whole organs were dissociated through 70 or 100 μm strainers into complete RPMI to generate single cell suspensions. The cell suspensions were then RBC lysed in ACK buffer, washed in complete RPMI and counted on Countess 3 (Invitrogen) to determine total viable cell number. The samples were plated on U-bottom 96-well plates at 3 × 10^6^ cells/well in complete RPMI.

For staining of lymph nodes, whole organs were placed into Click’s EHAA Medium (FUJIFILM Irvine Scientific). The samples were minced with needles and incubated in the presence of 0.1 mg/mL Collagenase D (Roche) and 0.02 mg/mL DNase I (Worthington Biochemical) with periodic resuspension. After 1 h incubation at 37°C, the enzymes were quenched by the Click’s EHAA media containing 5 mM EDTA and 2.5% FBS. The cells were filtered through strainer, washed, and counted on Countess 3 (Invitrogen) to determine total viable cell number.

Muscle tissue of the quadricep, containing the site of injection (SOI), was excised and processed into a single-cell suspension with the Skeletal Muscle Dissociation Kit (SMDK) (Miltenyi) and GentleMACS Octo with heaters (Miltenyi) according to the manufacturer’s recommendations. In short, excised tissue was minced, added to C-tubes (Miltenyi) with SMDK enzymes in DMEM, processed on standard program mr_SMDK_1 (3 min at +60 RPM, 9 min at -30 RPM using the heating function, 6 cycles of 5 sec at ± 360 RPM, 12 min at -30 RPM using the heating function, 3 cycles of 5 sec at ± 360 RPM) on the GentleMACS, filtered, and washed. The cells were counted and adjusted to 2 x 10^6^ live cells/well prior to staining.

### 2.5. Flow Cytometry

Cell suspensions were plated onto the U-bottom 96-well plate and incubated with anti-CD16/32 to block Fc receptors. To identify OVA-specific CD8+ T cells in blood and spleens by tetramer staining, the biotinylated Flex-T™ Biotin H-2 K(b) OVA Monomer (SIINFEKL) (BioLegend) was tetramerized with streptavidin conjugated to APC or BV421 (BioLegend) and added to the cells in the presence of CD8α (53-6.7, BioLegend) at 37°C for 30 min. Cells were washed with Flow Staining Buffer (Cytek) and stained with the viability dye (Ghost Dye Red 780 Fixable Viability Dye, Cell Signaling Technology), CD19 (6D5, BioLegend), CD44 (IM-7, Tonbo), CD127 (A7R34, Tonbo), KLRG1 (2F1/KLRG1, BioLegend) for 20 min at RT. Samples were washed, fixed for 30 min in Fixation Buffer (Tonbo), washed with Flow Staining Buffer, and permeabilized for five min with 1X Flow Cytometry Perm Buffer (Tonbo). After permeabilization, the samples were stained intracellularly with Granzyme B (QA16A02, BioLegend) for one h RT and resuspended in Flow Stain Buffer (Tonbo).

For the intracellular cytokine staining of the splenocytes, the cells were stimulated *ex vivo* with the OVA peptide pool (Peptivator Ovalbumin, Miltenyi), in the presence of Brefeldin A Solution (1000X) (Tonbo), anti-CD49d and anti-CD28 in complete RPMI media for 5 h at 37°C. Following incubation, cells were Fc blocked and surface stained with viability dye (Ghost Dye Red 780 Fixable Viability Dye, Cell Signaling Technology), CD19 (6d5, BioLegend), CD4 (GK1.5, BioLegend), CD8α (53-6.7, BioLegend), CD44 (IM7, Tonbo). Following surface staining, stimulated cells were fixed, permeabilized, and stained intracellularly with TNF-α (MP6-XT22, BioLegend), IL-2 (JESS-5H4, Tonbo), and IFN-γ (XMG1.2, Tonbo) for 30 min at RT, washed and resuspended in Flow Stain Buffer (Tonbo).

To analyze the immune subsets in muscle (SOI) and dLN, the cells were adjusted to 2×10^6^ cells/well and stained with the following combination of fluorochrome-conjugated mAbs: CD45 (30-F11), I-A/I-E (M5/114.15.2), CD11c (N418), CD8α (53-6.7), Ly6G (1A8), Ly6c (HkK1.4), B220 (RA3-6B2), CD11b (M1/70), CD3 (17A2), F4/80 (BM8) (Biolegend) and Ghost Dye Red780 (Tonbo) for live/dead discrimination. Staining was performed in the presence of CD16/32 (93, BioLegend) and True-Stain Monocyte Blocker™ (BioLegend) to block nonspecific binding of mAbs to Fc receptors or Myeloid CD64, respectively.

All samples were analyzed by flow cytometry using a Cytek Northern Lights 3-laser (VBR) spectral flow cytometer and analyzed using FlowJo software (BD Biosciences).

### 2.6. ELISpot

Spleens were harvested and single-cell suspensions were prepared by passing the organs through 70-μm cell strainers (Falcon) into RPMI 1640 (Gibco) supplemented with 10% HI fetal bovine serum and 1× penicillin/streptomycin (Gibco) (RPMI-10). Cells were pelleted by centrifugation (450 × g), resuspended in red blood cell lysis solution (Sigma), incubated at RT for 10 min, pelleted (450 × g), and then resuspended in RPMI-10 at a concentration of 5 × 106 cells/mL. Mouse IFN-γ Immunospot ELISpot kits (Cellular Technology Ltd.) were used per the manufacturer’s instructions. Splenocytes stimulated with an Hcp1 peptide pool or medium only were added to the plates at a concentration of 2.5 × 10^5^ cells/well and then incubated for 48 h at 37°C under an atmosphere of 5% CO_2_. The ELISpot plates were processed and developed per the manufacturer’s instructions. Plates were imaged using an ImmunoSpot S1 analyzer (Cellular Technology Ltd.). IFN-γ-secreting T cells were quantitated using the ImmunoSpot v5.1 professional DC smart count software (Cellular Technology Ltd.).

### 2.7. Statistical Analysis

GraphPad Prism (version 10.4.2, GraphPad) or Excel (Microsoft) were used for all statistical analyses. Figure legends detail the number of experimental replicates and n-values. Data presented are mean ± SD, mean ± SEM or Geometric Mean Ratio ± 95%CI. Significance was defined using unpaired t-test with Welch’s correction or the GMR analysis. *p* ≤ 0.05 were considered significant.

## 3. Results

### 3.1. Administration of OVA antigen formulated using ALTA^®^ platform elicits robust and durable antibody production measured by anti-OVA IgG1

ALTA^®^ particles were generated by formulating the model antigen OVA with excipients, spray-dried, and the resulting microparticles were then coated with aluminum oxide (Al_2_O_3_) using 50 cycles of ALD. To characterize the humoral response to OVA delivered within ALTA^®^ particles in comparison with the OVA formulated with classical adjuvants, C57BL6 mice were immunized i.m. with matching doses of antigen, and sera were collected over the course of 12 - 14 weeks to measure the OVA-specific IgG1 antibody titers by ELISA. After single immunizations with ALTA^®^delivering either 62.5, 250, or 1000 ng OVA, the antigen-specific IgG1 titers were at the background level at week 1 and peaked around weeks 2 and 4 post priming administration (**Figure 1A-D**, Supplementary Figure 1C-D). IgG1 titer levels were maintained over 12-14 weeks with less than one-log reduction compared with the peak titer. At the lowest OVA dose tested (62.5 ng), the anti-OVA IgG1 titer was higher after immunization with ALTA^®^ than when adjuvanted with Alhydrogel^®^, AddaVax™, or Alhydrogel^®^+CpG ODN 1018 (**Figure 1A**). The rate of seroconversion (arbitrarily defined as 2 log-fold increase in anti-OVA IgG1 titer relative to diluent control) was 7/10 for ALTA^®^, 1/10 for Alhydrogel^®^, 1/10 for AddaVax™, 0/10 for Alhydrogel^®^+CpG ODN 1018 at week 6 post prime. A further analysis was conducted of geometric mean ratios (GMR) between ALTA^®^ and the traditional liquid adjuvants. Although the 95% confidence interval (CI) was large for the ALTA^®^ group, the GMR analysis demonstrated that the liquid adjuvant formulations resulted in substantially reduced IgG1 titers (**Figure 1B**). In a separate study, the IgG1 production by ALTA^®^ was compared to poly(I:C)+anti-CD40 agonistic antibody. At 62.5 ng OVA, anti-OVA IgG1 was produced only after ALTA^®^ administration (Supplementary Figure 1A, B). At the 250 ng OVA dose, the IgG1 titers elicited by ALTA^®^ immunization were the highest among all tested groups (**Figure 1C**, D). At the 1000 ng OVA dose, the IgG1 titers were higher in animals treated with ALTA^®^ compared against AddaVax™ and the titers were similar between ALTA^®^ and Alhydrogel +/- CpG 1018 (Supplementary Figure 1C, D). Overall, a single administration of ALTA^®^ achieved superior IgG1 titers across time points at low antigen doses (62.5 and 250 ng OVA) and elicited similar IgG1 titers at high antigen doses (1000 ng OVA) when compared against the traditional adjuvanted formulations using Alhydrogel^®^, AddaVax™, Alhydrogel^®^+CpG ODN 1018, or poly(I:C) + anti-CD40.

**Figure 1.**
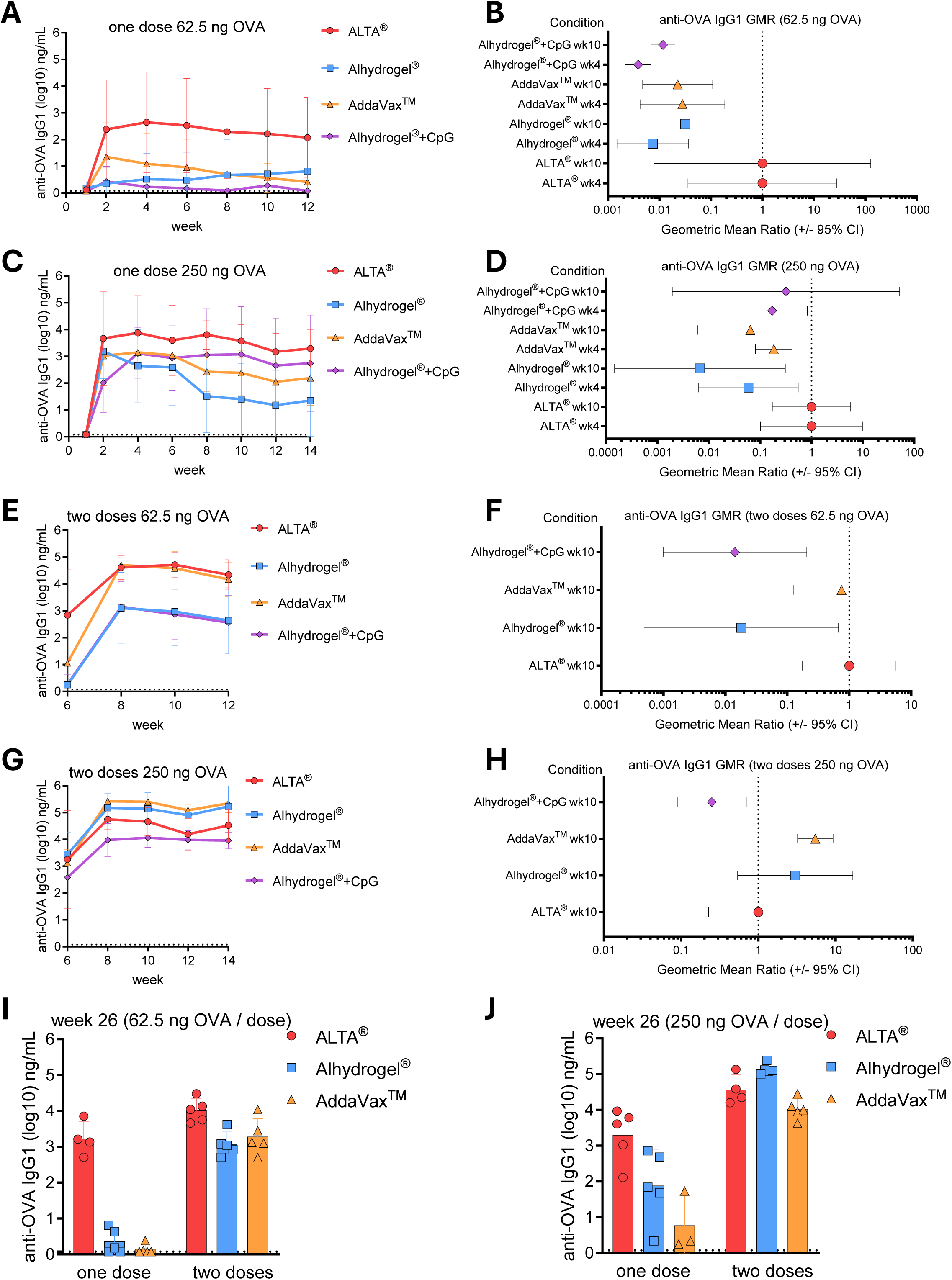
Administration of OVA antigen formulated using ALTA^®^ platform elicits robust and durable antibody production measured by anti-OVA IgG1. C57BL6 mice were injected i.m. with 62.5 or 250 ng of OVA antigen delivered in ALTA^®^ platform or with adjuvants, including Alhydrogel^®^, AddaVax™, Alhydrogel^®^+CpG ODN 1018. Where indicated, the mice received boost doses at week 6. Shown are the anti-OVA IgG1 titers measured by ELISA over the course of 12 - 14 weeks (A-H) and at week 26 (I, J). (A) Anti-OVA IgG1 titers (log10) over 12 weeks and (B) the geometric mean titers ratio (GMR) at weeks 4 (n=9-10) and week 10 (n=5) (62.5 ng OVA). (C) Anti-OVA IgG1 titers (log10) over 14 weeks and (D) the GMR at weeks 4 (n=10) and week 10 (n=5) (250 ng OVA). (E) Anti-OVA IgG1 titers (log10) before and after week 6 boost and (F) the GMR at week 4 post boost (n=4-5) (62.5 ng OVA/dose). (G) Anti-OVA IgG1 titers (log10) before and after week 6 boost and (H) the GMR at week 4 post boost (n=5) (250 ng OVA/dose). (I, J) Anti-OVA IgG1 (log10) at week 26 post one and two doses of 62.5 ng OVA/dose (n=4-5) (I) or 250 ng OVA/dose (n=3-5) (J). Shown are representative data from one of two (A-H: ALTA®, Alhydrogel^®^, AddaVax™ groups) or one experiment (A-H: Alhydrogel^®^+CpG group. I, J: all groups). Mean ± SD (A, C, E, G, I, J) and Geometric Mean Ratio ± 95% CI (B, D, F, H).

Because the traditional liquid subunit vaccine formulations typically require multiple doses to achieve desired magnitude and durability of the antibody titers, the IgG1 titers after two doses of ALTA^®^ were compared to two doses of liquid adjuvants. Mice were boosted at week 6 post prime dose with antigen doses matching prime administrations, and the antigen-specific IgG1 titers were measured bi-weekly until week 12-14. As expected, the second dose of ALTA^®^ delivering 62.5, 250 or 1000 ng resulted in increased IgG1 titers (**Figure 1E**, G, Supplementary Figure 1E). After two doses of 62.5 ng OVA, the IgG1 titers after ALTA^®^ were higher than Alhydrogel^®^ +/- CpG ODN 1018 and were similar to AddaVax™ (**Figure 1** E, F). At two doses of 250 ng OVA, the titers after ALTA^®^ were higher than Alhydrogel^®^ +/- CpG ODN 1018, similar to Alhydrogel^®^, and lower than AddaVax™ (**Figure 1G**, H). All immunizations with two doses of 1000 ng OVA achieved high IgG1 values (10^4^-10^6^ ng/mL) (Supplementary Figure 1E, F).

The durability of the humoral response is an important measure of vaccine efficacy^34^. To elicit a long-lasting response, most vaccines require multiple administrations of relatively high doses of the antigen along with an adjuvant. In a separate study, the durability of the humoral response to ALTA^®^ was measured and compared to the matching antigen doses of formulations containing Alhydrogel^®^ or AddaVax™ (**Figure 1I**, J). At week 26 post prime, all ALTA^®^immunized mice maintained elevated antigen-specific IgG1 titers > 2log_10_. In contrast, no animals had antigen-specific IgG1 titers > 2log_10_ after immunizations with single 62.5 ng OVA dose adjuvanted with either Alhydrogel^®^ or AddaVax™, and additional booster doses were required to achieve seroconversion in those animal groups. Two doses of OVA adjuvanted with either Alhydrogel^®^ or AddaVax™ elicited a similar response to a single dose of ALTA^®^ (**Figure 1I**). At a single 250 ng OVA dose, the highest titers were observed in the ALTA^®^ vaccinated mice (**Figure 1J**). At two doses, the IgG1 titers were high (>3log_10_) in all groups.

Taken together, these data show high levels of anti-OVA IgG1 antibody production were achieved after immunizations with one and two doses of ALTA^®^ containing OVA. After a single administration and at a low antigen dose, ALTA^®^ immunization elicited the highest OVA-specific IgG1 titers among all tested groups. Although the differences between the groups became less apparent at higher doses, antibody titers from ALTA^®^ vaccinated animals were similar when compared against animals vaccinated with most liquid adjuvants tested. Furthermore, antibody titers in ALTA^®^-vaccinated animals persisted for 26 weeks, suggesting that ALTA^®^ vaccines provide a durable humoral response.

### 3.2. Administration of OVA antigen formulated using ALTA^®^ platform elicits antibody subtypes that align with a balanced Th1/Th2 response

The addition of adjuvants to vaccine antigens can direct the immune response towards different functional states such as Th1, Th2, or other. Generally, adjuvants containing aluminum (Alhydrogel^®^ or Adju-Phos) elicit a Th2-skewed immune response characterized by high IgG1 production^14^. Other adjuvants, for example squalene oil-in-water emulsions, such as MF59 or AddaVax™, elicit both humoral (Th2) and cellular (Th1) immune responses ^24^. The addition of the TLR9 agonist, CpG oligonucleotide, to Alhydrogel^®^ is used to induce Th1 responses that would otherwise not occur with Alhydrogel^®^ alone^35^.

To elucidate whether the humoral response to ALTA^®^ is Th1 or Th2 skewed, C57BL6 mice were immunized as described above and the sera were analyzed for anti-OVA IgG2c antibody levels by ELISA (**Figure 2** and Supplementary Figure 1G-J)^36^. IgG2c was detected in sera two weeks post immunizations with all ALTA^®^ OVA doses tested, including 62.5, 250 and 1000 ng. At 62.5 ng single OVA dose, ALTA^®^ was the only treatment that elicited the IgG2c titers above the assay limit of detection (LoD) from two to ten weeks (**Figure 2A**). The IgG2c titers after ALTA^®^ OVA were significantly higher than after Alhydrogel^®^ OVA or AddaVax™ OVA immunizations at all doses and regiments tested (**Figure 2A-H**; Supplementary Figure 1G-J). The only condition that resulted in higher IgG2c titer values above the ALTA^®^ treated groups were animals immunized with a combination of CpG ODN 1018 and Alhydrogel^®^ adjuvants at the 250 or 1000 ng OVA doses (**Figure 2C** - D, G - H, Supplementary Figure 1G-J).

**Figure 2.**
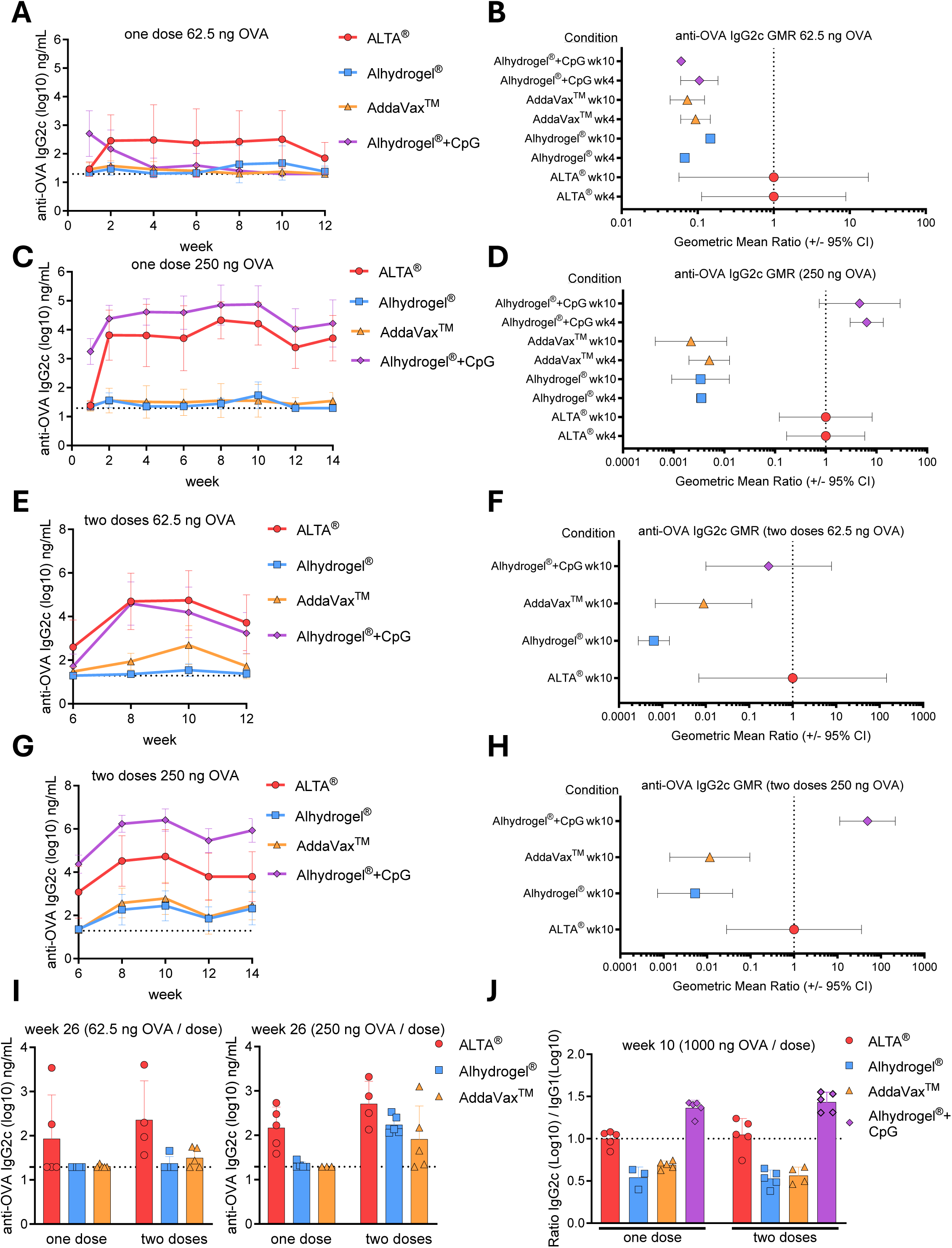
Administration of antigen formulated using ALTA^®^ platform elicits antibody subtypes that align with a balanced Th1/Th2 response. C57BL6 mice were injected i.m. with 62.5 or 250 ng of OVA antigen delivered in ALTA^®^ platform or with adjuvants, including Alhydrogel^®^, AddaVax™, Alhydrogel^®^+ CpG ODN 1018. Where indicated, the mice were boosted at week 6. Shown are the anti-OVA IgG2c titers measured by ELISA over the course of 12-14 weeks (**A**-**H**) and at week 26 (**I**). (**A**) Anti-OVA IgG2c titers (log10) over 12 weeks and (**B**) the GMR at week 4 (n=9-10) and week 10 (n=5) (62.5 ng OVA). (**C**) Anti-OVA IgG2c titers (log10) over 12 weeks and (**D**) the GMR at week 4 (n=5) and week 10 (n=5) (250 ng OVA). (**E**) Anti-OVA IgG2c titers (log10) before and after boost at week 6 and (**F**) the GMR at week 4 post boost (n=4-5) (62.5 ng OVA/dose). (**G**) Anti-OVA IgG2c titers (log10) before and after boost and (**H**) the GMR at week 4 post boost (n=5) (250 ng OVA/dose). (**I, J**) Anti-OVA IgG2c (log10) at week 26 post one or two doses of 62.5 ng OVA/dose (n=4-5) or 250 ng OVA/dose (n=3-5). (**J**) The ratio of IgG2c (log10) to IgG1 (log10) at week 10 (1000 ng OVA/dose, n=3-5/group). Shown are representative data from one of two (**A**-**H**: ALTA^®^, Alhydrogel^®^, AddaVax™ groups) or one experiment (**A**-**H**: Alhydrogel^®^+CpG group. **I**, **J**: all groups). Mean ± SD (**A**, **C**, **E**, **G**, **I**, **J**) and Geometric Mean Ratio ± 95% CI (**B**, **D**, **F**, **H**).

In a separate study, the durability of IgG2c production post ALTA^®^ immunization in comparison to Alhydrogel^®^ and AddaVax™ formulations was measured at week 26 post prime (**Figure 2I**). In contrast to liquid adjuvants, the IgG2c was present in all ALTA^®^ immunized groups.

To determine whether immunization with ALTA^®^ skews the humoral response to Th1 or Th2, the ratio of the anti-OVA IgG2c and IgG1 was calculated (**Figure 2J**) ^37,38^. Here, the 1000 ng OVA dose was chosen for analysis to ensure that all mice had IgG2c and IgG1 values above the LoD. At week 10, the IgG2c to IgG1 ratio was below 1 for the Alhydrogel^®^ and AddaVax™ groups, demonstrating a Th2 bias. The ratio was above 1 for the Alhydrogel^®^ + CpG 1018 group demonstrating the Th1 skewing after the addition of CpG 1018. The ALTA^®^ immunized group was the only treatment that achieved the ratio closest to 1, suggesting a balanced immune response post this vaccination.

### 3.3. Administration of antigen formulated using ALTA^®^ platform elicits a more robust antigen (OVA)-specific CD8+ T cell response than formulations containing Alhydrogel^®^, AddaVax™, Alhydrogel^®^+CpG ODN 1018

Generally, the serum antibody responses strongly correlate with protection and serve as a primary read-out for the characterization of vaccine efficacy^39,40^. However, cell-mediated immunity is also an important measure of vaccine efficacy^41,42^, especially in the context of vaccines against intracellular pathogens or cancers^43–45^. The above data showed that vaccination with ALTA^®^ containing OVA elicited a robust and durable humoral immunity characterized by the anti-OVA IgG1 and IgG2c production. Whether ALTA^®^ elicited a T cell response and how it compared to the liquid adjuvants remained to be determined. To measure the CD8+ T cell response, C57BL6 mice were immunized as described above and whole blood was analyzed at week 1, 2, 4, 8 and10 (**Figure 3**). OVA-specific CD8+ T cells were identified by the Major Histocompatibility Complex (MHC) class I tetramer staining and flow-cytometry (Supplementary Figure 2A). The frequency of the OVA-specific CD8+ T cells was low at all time points and all doses after immunizations with OVA adjuvanted with Alhydrogel^®^, AddaVax™, or Alhydrogel^®^+CpG ODN 1018 (**Figure 3B**, C, Supplementary Figure 2B, C). Among those groups, the highest mean frequency (>0.5%) of the OVA-specific CD8+ T cells was detected at week 1 post immunization with 1000 ng OVA + Alhydrogel^®^ (Supplementary Figure 2C). In contrast, the OVA-specific CD8+ T cells were easily detectable in the blood of the ALTA^®^-vaccinated mice at all time points past week 1 and all doses tested. After immunization with ALTA^®^, the OVA-specific CD8+ T cell responses peaked at week 2 and then contracted. The percent of the OVA-specific CD8+ T cell responses in blood was higher after ALTA^®^ immunization than liquid adjuvants at all time points beyond week 2. In a separate experiment, the OVA-specific CD8+ T cell responses were measured after immunizations with ALTA^®^ or OVA formulated with a combination of the TLR3 agonist poly(I:C) and anti-CD40 agonistic antibody (Supplementary Figure 2D). At this antigen dose (62.5 ng OVA), only ALTA^®^ treatment elicited an appreciable OVA-specific CD8+ T cell response in blood.

**Figure 3.**
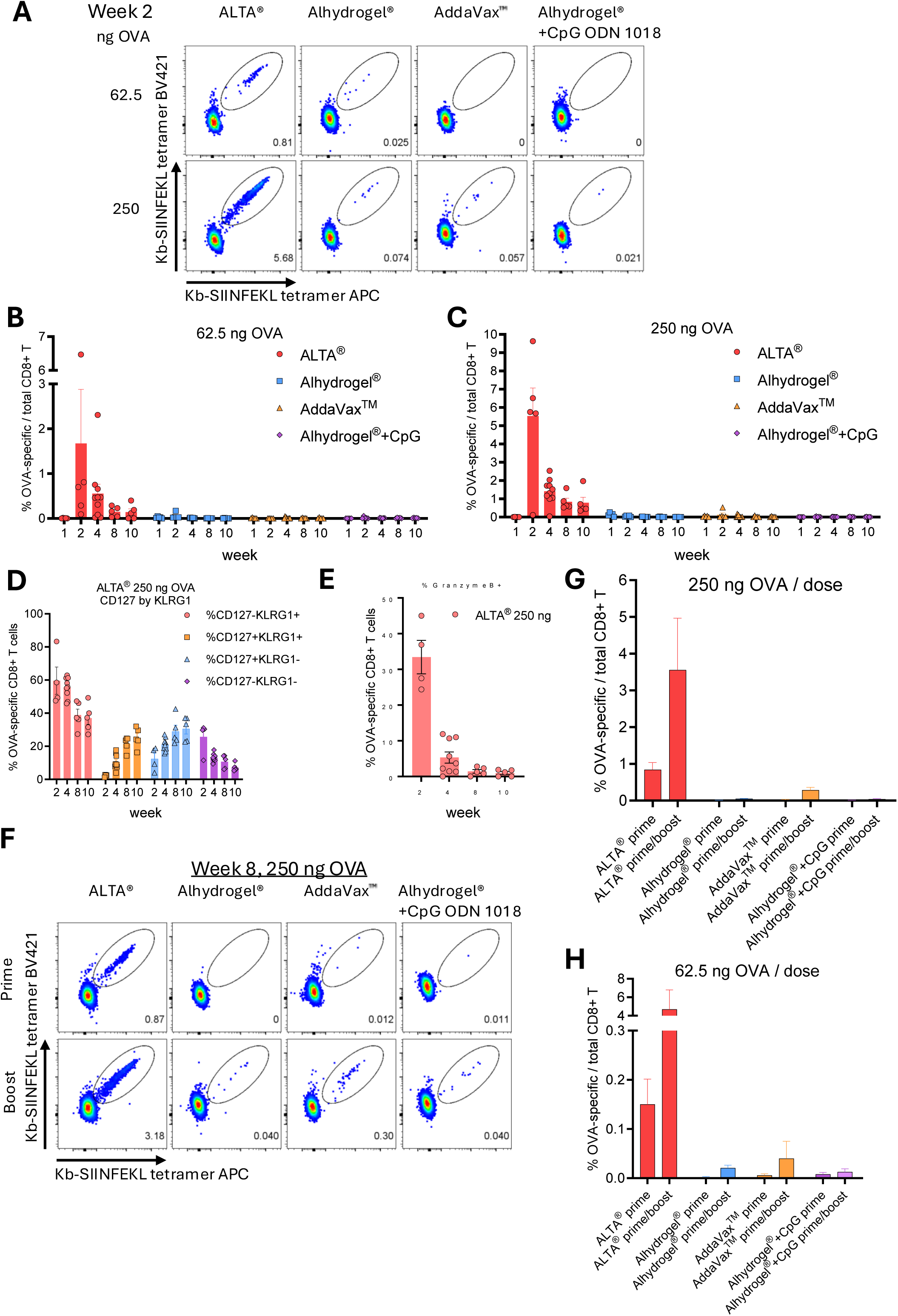
Administration of antigen formulated using ALTA^®^ platform elicits a more robust antigen (OVA)-specific CD8+ T cell response than Alhydrogel^®^, AddaVax™, Alhydrogel^®^+CpG ODN 1018. C57BL6 mice were i.m. immunized with 62.5 or 250 ng of OVA antigen delivered in ALTA^®^ or with Alhydrogel^®^, AddaVax™, Alhydrogel^®^+CpG ODN 1018. Blood was collected at the indicated time points after prime (**A**-**E**) or boost (week 6) (**F**-**H**) and analyzed for presence and phenotype of the OVA-specific CD8+ T cells by flow cytometry. (**A**) Representative dot plots depicting the percentage of OVA-specific CD8+ T cells of total CD8+ T cells (gated as H-2K^b^-SIINFEKL tetramer APC+ H-2K^b^-SIINFEKL tetramer BV421+ in CD8α+CD19- live lymphocytes) at week 2 post-vaccination. (**B, C**) The frequency of OVA-specific CD8+ T cells in blood at week 1, 2, 4, 8, and 10 post vaccinations with 62.5 (**B**) or 250 ng (**C**) OVA. The percent of CD127 (IL-7Rα) and/or KLRG1-expressing (**D**) and Granzyme B+ (**E**) within OVA-specific CD8+ T cells after ALTA^®^ vaccination (250 ng OVA). (**F**) Representative dot plots depicting the percentage of OVA-specific CD8+ T cells (gated as H-2K^b^-SIINFEKL tetramer APC+ H-2K^b^-SIINFEKL tetramer BV421+ in CD8α+CD19- live lymphocytes) at week 8 post prime (top row) and two weeks after boost (bottom row). The frequency of OVA-specific CD8+ T cells at week 8 after prime and two weeks post week 6 boost with 250 ng (**G**) and 62.5 ng OVA (**H**). Shown are the representative data from one of two (**A**-**E**: ALTA^®^ and Alhydrogel® groups) or one experiment (**A**-**D**: AddaVax™ and Alhydrogel^®^+CpG ODN 1018 groups. **F-H**: all groups). N=4-10/group. Mean ± SEM.

The phenotype of the OVA-specific CD8+ T cells was measured by the surface staining for the effector cell marker KLRG1 and the memory cell marker CD127 (IL-7Rα) (**Figure 3D**)^46^. At the peak response (week 2) to ALTA^®^ immunization, most CD8+ T cell responses displayed an effector (KLRG1+CD127-) and early effector KLRG1-CD127-) cell phenotype. A transitioning to the memory cell phenotype (CD127+) was observed at the later time points post-ALTA^®^. At weeks 8 and 10 the phenotype of the ALTA^®^-elicited CD8+ T cell responses was distributed between CD127+ (memory) and KLRG1+ (late effector) subsets. Additionally, the cytotoxic potential of the OVA-specific CD8+ T cell responses was measured by the intracellular staining for Granzyme B^47^. Consistent with peak overall percentages observed at week 2, the highest percent of the Granzyme B+ CD8+ T cells was detected at week 2 post ALTA^®^ (**Figure 3E**). These data showed, a single immunization with ALTA^®^ OVA elicited a more robust OVA-specific CD8+ T cell response in blood than OVA adjuvated with Alhydrogel^®^, AddaVax™, Alhydrogel^®^+CpG ODN 1018, and poly(I:C)+anti-CD40.

To measure the T cell response to the boost dose of ALTA^®^ in comparison with the liquid adjuvants, the mice were injected with the matching second doses to the prime vaccines at week 6 post prime. As expected, the frequency of the OVA-specific CD8+ T cell responses in blood increased after the boost doses of ALTA^®^ or liquid adjuvanted OVA (**Figure 3F-H**, Supplementary Figure 2E, F). At all doses tested, the frequency of the OVA-specific CD8+ T cell responses was higher after two doses of ALTA^®^ than liquid adjuvanted OVA. Moreover, the OVA-specific CD8+ T cell responses were detected at a higher frequency after a single administration of ALTA^®^ than two doses of the liquid adjuvants. The only liquid adjuvanted vaccine capable of achieving a comparable response to a single dose of ALTA^®^ was AddaVax™ given twice at a 1000 ng OVA per dose (Supplementary Figure 2F). Collectively, these data show that the ALTA^®^ platform elicits a strong cell-mediated immunity (from one or two administrations) as measured by the antigen-specific CD8+ T cell responses in blood and at levels greater than observed for the liquid vaccine formulations tested.

### 3.4. The frequency and numbers of the total and cytokine-producing OVA-specific CD8+ T cells in the spleens are increased after ALTA^®^ immunization

The numbers and functionality of the antigen-specific CD8+ T cell responses after immunizations with one or two doses of ALTA^®^ in comparison with the liquid adjuvanted OVA were measured in the secondary lymphoid organ, spleen. The mice were immunized, as described previously, and harvested at 14 weeks post prime (eight weeks post boost). First, the frequency and numbers of the OVA-specific CD8+ T cells were determined by the staining with the fluorescently labeled H-2K^b^ SIINFEKL tetramers and measured via flow-cytometry. In line with the frequencies of the OVA-specific CD8+ T cells in blood (Supplementary Figure 2C), both the frequency and numbers were higher after immunizations with ALTA^®^ than liquid adjuvanted OVA (**Figure 4A**, B). Among liquid vaccine regimens tested, only two doses of OVA adjuvanted with AddaVax™ elicited CD8+ T cell response in the spleen comparable to single dose of OVA-ALTA^®^. At 14 weeks post ALTA^®^ vaccine administration, the phenotype of the OVA-specific CD8+ T cells in spleen was distributed between the memory (CD127+) and effector (KLRG1+) cells (**Figure 4C**, D).

**Figure 4.**
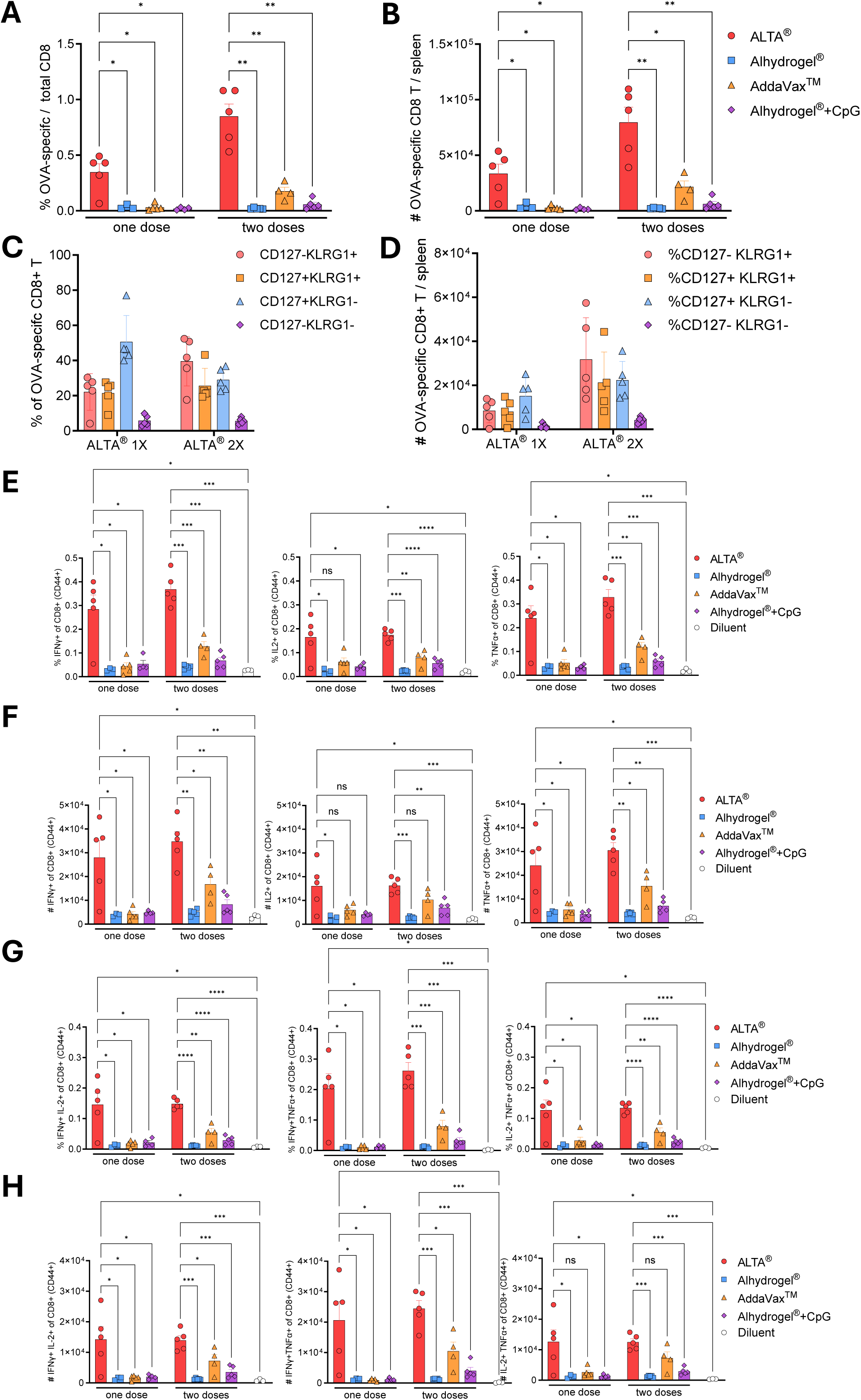
The frequency and numbers of the total and cytokine-producing OVA-specific CD8+ T cells in the spleens are increased after ALTA^®^ immunization. C57BL6 mice were i.m. immunized with 1000 ng of OVA delivered in ALTA^®^ or with Alhydrogel^®^, AddaVax™, Alhydrogel^®^+CpG ODN 1018. Indicated groups were boosted at week 6 post prime. (**A**-**D**) The spleens were collected at week 14 post prime and analyzed for presence and phenotype of the OVA-specific CD8+ T cells by flow cytometry (gated as H-2K^b^-SIINFEKL tetramer APC+ H-2K^b^-SIINFEKL tetramer BV421+ in CD44+CD8α+CD19- live lymphocytes). (**A**) The percentage of OVA-specific CD8+ T cells of total CD8+ T cells. (**B**) The total number of OVA-specific CD8+ T cells per spleen. (**C**) The percent and number (**D**) of CD127 (IL-7Rα) and/or KLRG1-expressing OVA-specific CD8+ T cells after ALTA^®^ OVA vaccination. (**E**-**H**) The splenocytes were re-stimulated with the OVA peptide pool for 5 h *ex vivo*, and the expression of IFN-γ, IL-2, and/or TNF-α was measured by intracellular cytokine staining. (**E, F**) The frequency (**E**) and number per spleen (**F**) of the IFN-γ, IL-2, or TNF-α -expressing CD8+ T cells (gated as CD44+CD8α+CD19- live lymphocytes). (**G**, **H**) The frequency (**G**) and number per spleen (**H**) of the CD8+ T cells expressing two cytokines (IFN-γ+IL-2+, IFN-γ+TNF-α+, IL-2+TNF-α+). N=3-5/ group. Mean ± SEM. Unpaired t test with Welch’s correction. *p ≤ 0.05, **p ≤ 0.01, ***p ≤ 0.001, ****p ≤ 0.0001.

To measure functionality of the OVA-specific CD8+ T cell responses generated by immunizations, splenocytes were restimulated *ex vivo* with an OVA-derived peptide pool and analyzed for the expression of the cytokines (**Figure 4E-H**). The frequency and numbers of the IFN-γ, TNF-α, and IL-2 expressing CD8+ T cells were higher in the spleens of the ALTA^®^ immunized mice compared to the diluent injected control mice (**Figure 4E-F**). In addition, the CD8+ T cells expressing two cytokines at the same time (gated as IFN-γ+IL-2+, IFN-γ+TNF-α+, IL-2+TNF-α+) were also present in the spleens of the ALTA^®^ immunized mice (**Figure 4G**, H). In comparison to the groups immunized with the liquid adjuvanted OVA, the frequency and numbers of the total IFN-γ+, TNF-α+, or IL-2+ and polyfunctional IFN-γ+IL-2+, IFN-γ+TNF-α+, or IL-2+TNF-α+ CD8+ T cells were generally higher after ALTA^®^ immunization. Among the liquid adjuvanted groups, the most robust cytokine expression was achieved after two doses of AddaVax™ adjuvanted OVA; however, it was comparable or lower than one dose of ALTA^®^. In addition to being present and functional at week 14 post prime, the OVA-specific CD8+ T cell responses were also detectable as late as 34 weeks post ALTA^®^ OVA (250 ng) immunization (data not shown). Overall, one or two doses of ALTA^®^ elicited a more robust and long-lasting antigen-specific CD8+ T cell response in blood and spleen as compared to the liquid vaccine formulations tested.

To identify OVA-specific CD4+ T cells and measure their functionality after immunizations with ALTA^®^ versus liquid adjuvanted OVA, splenocytes were restimulated *ex vivo* with OVA-derived peptide pool in the presence of co-stimulatory molecules, CD28 and CD49d. Increased frequency and numbers of the total IFN-γ, IL-2, or TNF-α expressing CD4+ T cells were detected after ALTA^®^ vaccination as compared to the diluent control (Supplementary Figure 3A, B). The frequency and numbers of the cytokine-producing CD4+ T cells were relatively low after single administrations in all groups. After two doses, the CD4+ T cell response in the ALTA^®^ group was higher than in Alhydrogel^®^ and similar to the Alhydrogel^®^+CpG groups. Among all groups tested, two doses of vaccine adjuvanted with AddaVax™ achieved the highest CD4+ T cells with IL-2 and TNF-α expression. The percent and number of the polyfunctional CD4+ T cells (gated as IFN-γ+IL-2+, IFN-γ+TNF-α+, IL-2+TNF-α+) after ALTA^®^ immunization were greater than the percent and number observed in the diluent control group (Supplementary Figure 3C, D). The CD4+ T cell subsets producing two cytokines were more abundant after ALTA^®^ immunization than the liquid vaccine formulation containing Alhydrogel^®^. While the CD4+ T cell response was increased after the addition of CpG to Alhydrogel^®^, two doses were required to achieve a similar response compared to ALTA^®^ group. Consistent with increased IL-2 and TNF-α production, the IL-2+TNF-α+ subset was the most abundant after two doses of OVA adjuvanted with AddaVax™. Overall, these data show that immunization with ALTA^®^ vaccine formulation elicited a detectable and functional antigen-specific CD4+ T cell response.

### 3.5. ALTA^®^ platform is antigen and aluminum sparing

The data above demonstrated that immunization with OVA containing ALTA^®^ particles was efficient in eliciting robust humoral and cellular antigen-specific responses. A single administration with ALTA^®^ particles delivering low antigen doses (62.5 and 250 ng OVA) led to the production of higher antigen-specific antibody (IgG) titer levels than other vaccine formulations at equivalent doses. These data demonstrated immunization with ALTA^®^ required less OVA antigen to elicit seroconversion, suggesting an antigen-sparing capacity of ALTA^®^ (**Figure 5A**, B). To increase vaccine efficacy, most vaccines containing Alhydrogel^®^ require high doses of Alhydrogel^®^ that translates to higher amounts of elemental aluminum per dose^15^. ALTA^®^ particles are coated with nanoscopic alumina (Al_2_O_3_) layers and thereby employ significantly less aluminum (**Figure 5A**, B). For example, at 62.5 ng OVA dose, ALTA^®^ demonstrated an increased potency over the Alhydrogel^®^ +/- CpG formulations. In these conditions, the Alhydrogel^®^ formulations contained 3,125 ng Al^3+^, whereas the ALTA^®^ formulation contained ∼40-fold less Al^3+^ (79 ng Al^3+^). Therefore, in comparison to the traditional aluminum-containing adjuvant Alhydrogel^®^, ALTA^®^ provides an aluminum-sparing capacity with an improved immunogenicity.

**Figure 5.**
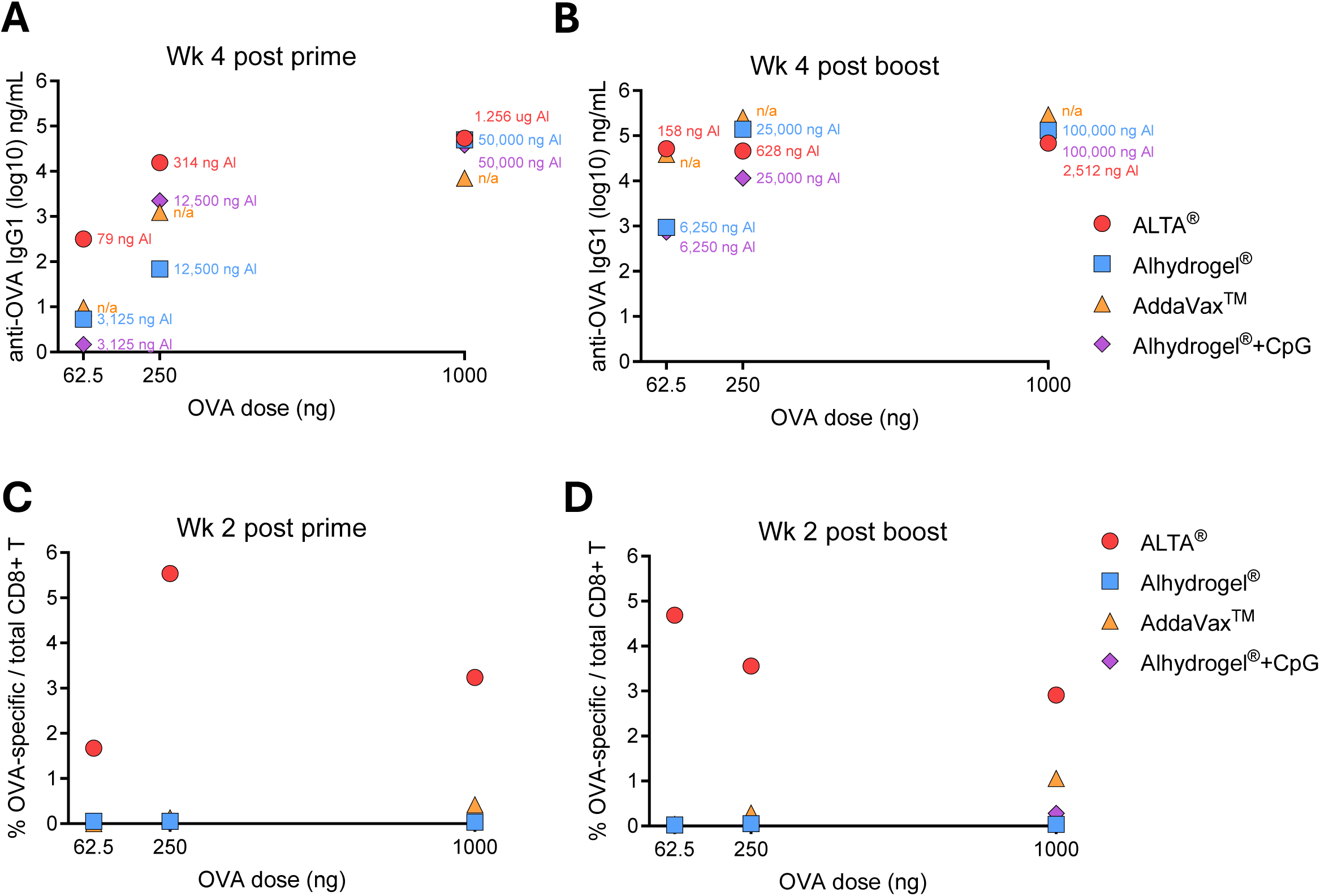
ALTA^®^ platform is antigen and aluminum sparing. C57BL6 mice were i.m. immunized with 62.5, 250 or 1000 ng of OVA antigen delivered in ALTA^®^ or with Alhydrogel^®^, AddaVax™, Alhydrogel®+CpG ODN 1018. Where indicated, the boost doses were administered at week 6 post prime. (**A**, **B**) Shown are the OVA doses (ng) (X axis) by the mean anti-OVA IgG1 (log10) titers at week 4 post prime (**A**) and week 4 post boost (Y axis). (**B**) The amount of aluminum (ng Al3+) delivered with each treatment is indicated as a color-coded number on the graph. The AddaVax™ group did not receive any aluminum and is labeled as “n/a” (not applicable). (**C**, **D**) Shown are the OVA doses (ng) (X axis) by the mean percent of the OVA-specific CD8+ T cells in blood at week 2 post prime (**C**) and week 2 post boost (**D**). N=4-5 /group.

At all antigen doses tested in this study, vaccine formulations adjuvanted with Alhydrogel^®^+CpG and AddaVax™ demonstrated a limited capacity to elicit OVA-specific CD8+ T cell responses in both blood and spleens (**Figure 5C**, D). To achieve an appreciable level of the T cell response, vaccine adjuvanted with AddaVax™ required two doses of 1000 ng OVA. In contrast, ALTA^®^ induced robust CD8+ T cell responses at all doses, once again demonstrating its potency and an antigen-sparing capacity (**Figure 5C**, D). Thus, in comparison to the established capacity for vaccines adjuvanted with AddaVax™ or Alhydrogel^®^+CpG to induce appreciable cell-mediated immune responses, ALTA^®^ vaccines generated more robust CD8+ T cell responses with less antigen, significantly lower aluminum amounts, and in fewer doses, demonstrating its superiority as a vaccine platform modality.

### 3.6. The immunogenicity of the ALTA^®^ platform is dependent on antigen containment within particles

Differences in the magnitude and quality of the immune responses to the antigen delivered with ALTA^®^ platform versus traditional adjuvants were observed. In the experiments described above, the antigens for all ALTA^®^ conditions were contained within the ALD-coated particles. It was hypothesized that this was a critical feature contributing to the immunogenicity of the ALTA^®^ platform and that a significant reduction in the immune response would be observed if the antigen was no longer contained within the particles. To test this, mice were immunized with placebo ALTA^®^ particles containing no antigen and supplemented with OVA outside the particle, in the diluent (ALTA^®^ placebo + OVA group). Additionally, to test whether integrity or intactness of the particles was also important for the immunogenicity of ALTA^®^, the particles (ALTA^®^ OVA or ALTA^®^ placebo) were intentionally broken through physical disruption and injected i.m. This physical disruption method resulted in roughly 86% of the particles broken with immediately available antigen. Similarly to prior results, immunization with the intact ALTA^®^ OVA elicited a robust and durable anti-OVA IgG1 response (**Figure 6A**). However, the antibody titers were substantially lower in the groups immunized with the placebo or broken ALTA^®^ particles. To measure the cellular immunity, these groups were harvested at week 22 and spleens were analyzed for OVA-specific CD8+ T-cell responses. Compared to ALTA^®^ OVA, the percentage and number of OVA-specific CD8+ T cells were significantly reduced in the spleens of mice immunized with the placebo or broken ALTA^®^ (**Figure 6B**, C). Therefore, the containment of the antigen inside the particles and the integrity of the ALTA^®^ particles are important to impart the superior immunogenicity of the ALTA^®^ platform.

**Figure 6.**
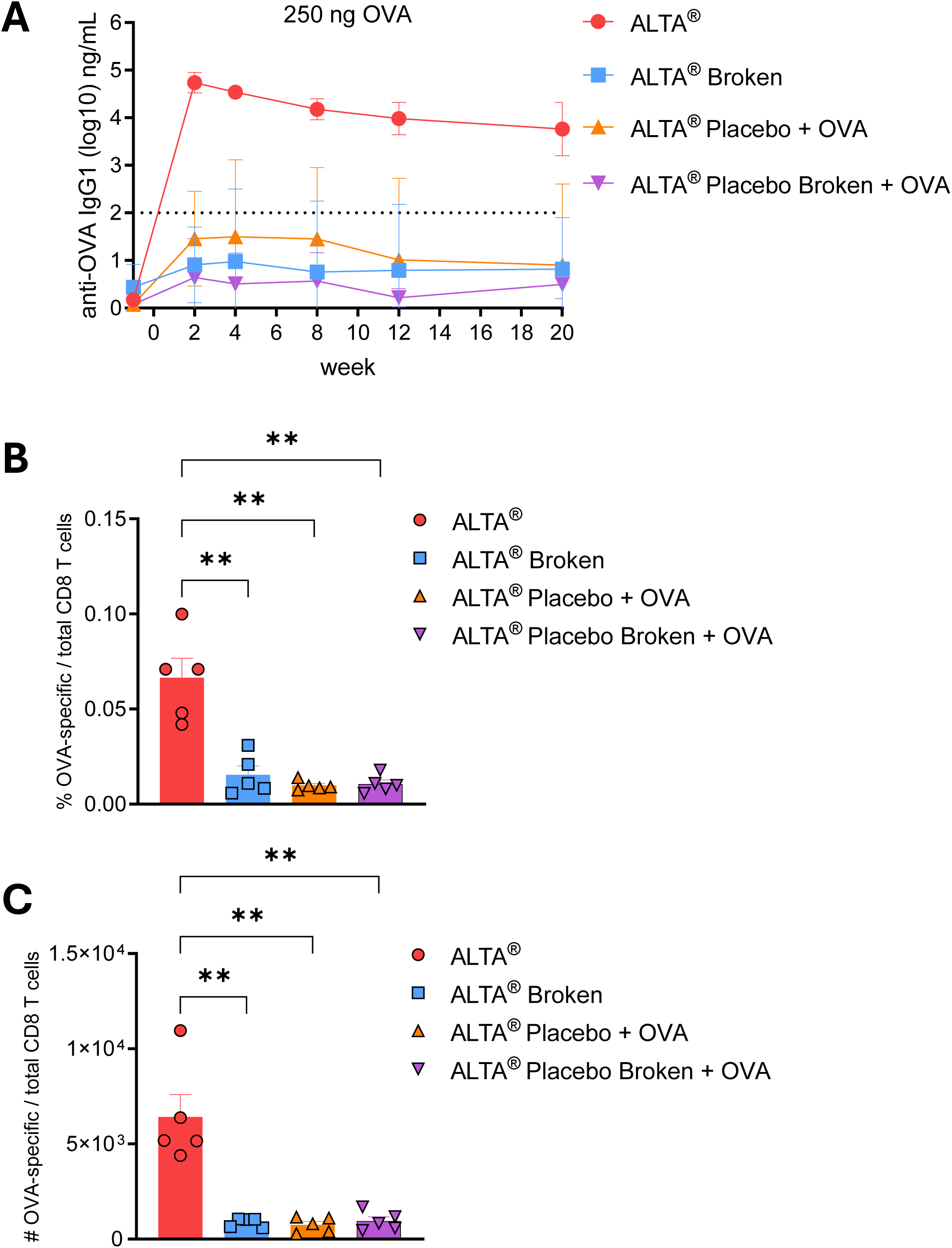
The immunogenicity of the ALTA^®^ platform is dependent on antigen containment within particles. C57BL6 mice were i.m. immunized with 250 ng of OVA delivered within ALTA^®^ particles (ALTA^®^ OVA), in diluent with ALTA® particles formulated without antigen (ALTA® placebo), broken ALTA® particles (ALTA^®^ broken), or in diluent with broken ALTA® placebo particles. (**A**) Anti-OVA IgG1 titers (log10) over 20 weeks. N=5. Mean ± SD. (**B**, **C**) Bar graphs depicting the percent (**B**) and number (**C**) of OVA-specific CD8+ T cells (gated H-2K^b^-SIINFEKL tetramer APC+ H-2K^b^-SIINFEKL tetramer BV421+ in CD8α+CD19- live lymphocytes) in the spleens at 22 weeks post immunizations. Shown are representative data from one of two or more experiments. N = 5 /group. Mean ± SEM. Unpaired t test with Welch’s correction. **p ≤ 0.01.

### 3.7. Immunization with ALTA^®^ particles recruits immune cells to the site of injection and increases antigen capture

To further characterize the immune response to ALTA^®^ formulated antigen and investigate mechanisms underlying ALTA^®^ platform immunogenicity, the site of injection (muscle) was assessed for the presence of major immune cell subsets. To this end, animals were given a single i.m. administration of ALTA^®^ particles containing OVA, ALTA^®^ placebo particles with OVA added to the diluent, liquid OVA, or diluent alone. The changes in the main immune cell subsets, including B and T cells, macrophages, neutrophils, Ly6C^hi^ monocytes, and dendritic cells (DCs), were assessed at days 1, 3, 14, and 42 following injections (Supplementary Figure 5A-G). Vaccination with ALTA^®^ particles, regardless of whether OVA was inside or in the diluent, resulted in higher numbers of innate immune cell subsets at the site of injection when compared to controls (Supplementary Figure 5E, F). These data demonstrate that ALTA^®^ particles are intrinsically immunostimulatory and may facilitate recruitment of immune cells to the site of injection.

Next, antigen positive cells were characterized at the site of injection using a fluorescent IVISense680 label that was conjugated to OVA. As expected, fluorescent OVA+ cells were detected in all groups except diluent (**Figure 7** and Supplementary Figure 5H-M). Compared to the liquid OVA group, the numbers and frequency of total OVA+ events were higher in both ALTA^®^ groups (**Figure 7A**, B). The number and frequency of OVA+ events was then determined within main APC subsets (**Figure 7C-F**, Supplementary Figure 5H-J). The peak number of OVA+ macrophages was seen one day following ALTA^®^ OVA administration and this peak was greater than the peaks observed across all other treatment groups through day 14 (**Figure 7C**). The numbers of OVA+ DC at the site of injection were trending higher for animals immunized with ALTA^®^ OVA compared to the other groups (**Figure 7D**). Interestingly, the numbers of OVA+ cells were higher after administration of OVA in the presence of ALTA^®^ placebo particles than in their absence (compare Placebo ALTA^®^ OVA *vs* OVA group). This result suggested that the ALTA^®^ particles have an intrinsic capacity to promote antigen capture at the site of injection, perhaps through the recruitment of innate immune cells. However, the containment of antigen inside versus outside of the particles elevated the numbers of the antigen-positive cells at the site of injection (**Figure 7A**). Collectively, these data suggest that ALTA^®^ particles recruit innate immune cells to the site of injection but that antigen uptake and/or maintenance at the site of injection is enhanced when antigen is contained within these particles, which correlates with an increased adaptive immunity (**Figure 6**).

**Figure 7.**
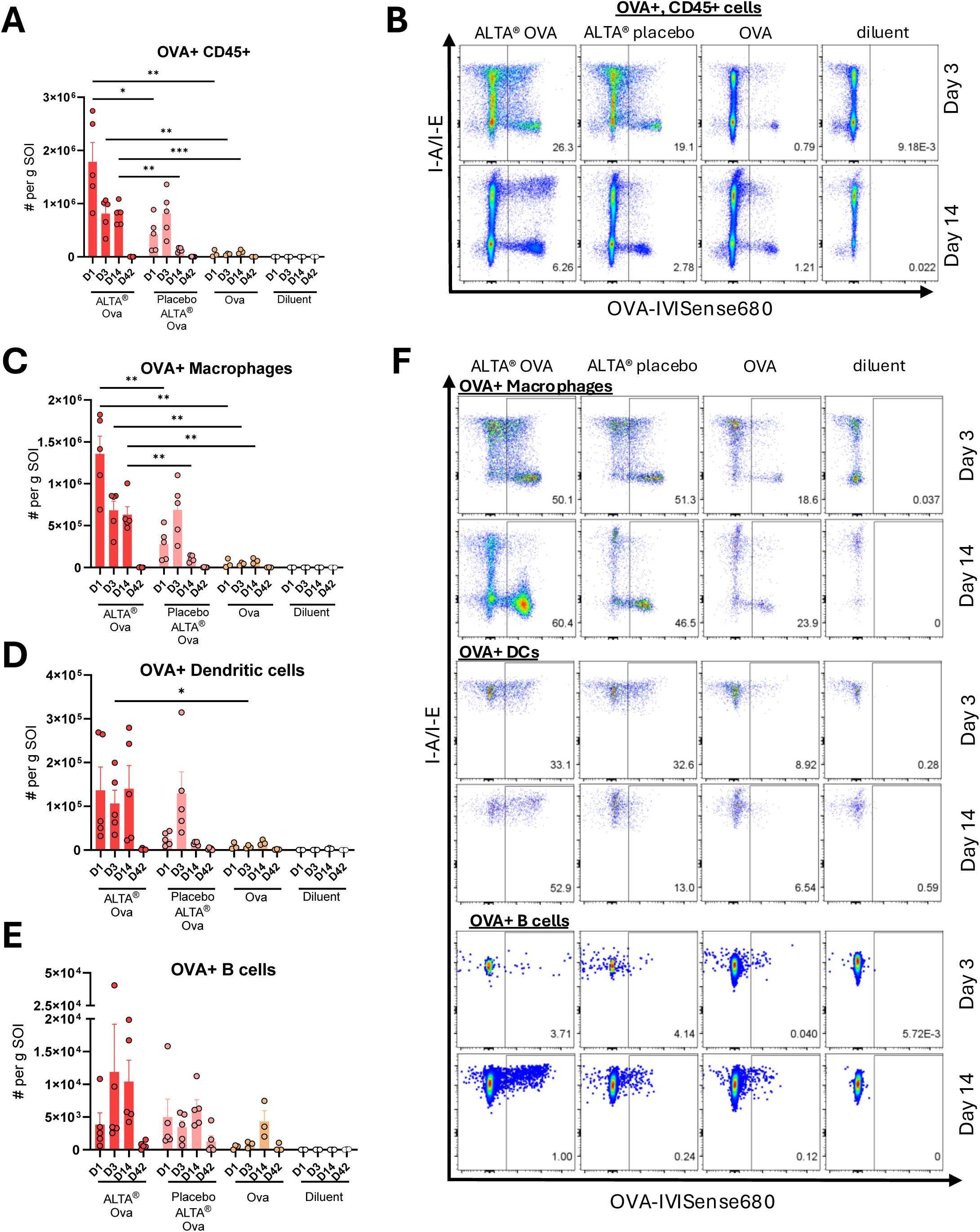
Immunization with ALTA^®^ particles recruits immune cells to the site of injection and increases antigen capture. C57BL6 mice were i.m. immunized with OVA-IVISense680 dye conjugate formulated using ALTA^®^ platform (ALTA^®^ OVA) or OVA-IVISense680 dye conjugate mixed in diluent with placebo ALTA^®^ particles formulated without antigen (Placebo ALTA^®^). Liquid control groups were injected with liquid OVA-IVISense680 or diluent. All groups except diluent control were administered 820 ng OVA dose. At days 1, 3, 7, and 42, muscle tissue containing site of injection was harvested and analyzed by flow cytometry. (**A**) Absolute cell number per gram muscle tissue of OVA-IVIS680+ CD45+ immune cells. Cell populations were gated on singlets, live cells, CD45+, OVA+. Absolute count of OVA+ CD45+ per gram muscle was determined as follows: [(%OVA+/100)*(%CD45+/100)*(live cell count)]/grams muscle mass. (**B**) Representative flow cytometry plots displaying percent of OVA+ of total CD45+ cells in injected muscle at days 3 and 14 post treatments. (**C**-**E**) Absolute cell number of OVA+ APC subset per gram injected muscle. Absolute number of OVA+ APCs per gram muscle were determined as follows: [(%OVA+/100)*(%APC/100)*(%CD45+/100)*(live cell count)]/grams muscle. Antigen-presenting cell populations were gated on singlets, live cells, CD45+, CD3- subsets ahead of the shown plots. (**C**) Macrophages were identified as CD45+, CD11b+, Ly6G-, Ly6c int/-, and F4/80+. (**D**) DCs were identified as CD45+, I-A/I-E+, CD11c^hi^. (**E**) B cells were identified as CD45+, I-A/I-E+, B220+. (**F**) Representative flow cytometry plots displaying percent of OVA+ of macrophages, DCs, B cells in injected muscle at days 3 and 14 post treatments. Shown are representative data from one of two experiments. N=3-5/group. Mean ± SEM. Unpaired t test with Welch’s correction (shown are comparisons between select groups (ALTA® OVA vs ALTA® Placebo; ALTA® OVA vs OVA) at matching time points). *p ≤ 0.05, **p ≤ 0.01, ***p ≤ 0.001.

### 3.8. Antigen containment within ALTA^®^ particles results in an increased number of antigen-positive cells in the dLN

Vaccine efficacy is dependent on the presence and processing of the antigen in the dLNs^14^. The iliac LN is the principal dLN post i.m. vaccination in the hind quadricep^48^. The presence of the main immune cell subsets at the iliac LN was interrogated by flow cytometry post single i.m. administration of ALTA^®^ OVA, ALTA^®^ placebo with OVA in diluent, liquid OVA, or diluent at days 3, 14 and 42 (Supplementary Figure 6A-G). The total number of hematopoietic cells as well as all individual subsets were generally higher post vaccination with ALTA^®^ OVA when compared to other groups. The most prevalent subsets post ALTA^®^ vaccination were T and B cells, followed by macrophages and DCs, followed by Ly6C^hi^ monocytes and finally neutrophils.

Transport of vaccine antigens to the dLNs and their retention for a sufficient duration are crucial for mounting a strong and durable immune response^49^. The number and frequency of OVA-IVISense680+ CD45+ cells were substantially higher at day 14 after immunization with ALTA^®^ OVA particles than other treatments (**Figure 8A**, B). These data show ALTA^®^ particles containing antigen facilitate delivery and/or maintenance of the antigen to and within the dLNs. The number and frequency of OVA+ cells were then determined within main APC subsets (**Figure 8C-F**, Supplementary Figure 6H-J). Most OVA+ cells in the ALTA^®^ immunized group were distributed between B cells and DCs, followed by macrophages. The highest number of OVA+ macrophages, DCs, and B cells was found at day 14 post injection of OVA formulated within ALTA^®^ platform (**Figure 8C-E**).

**Figure 8.**
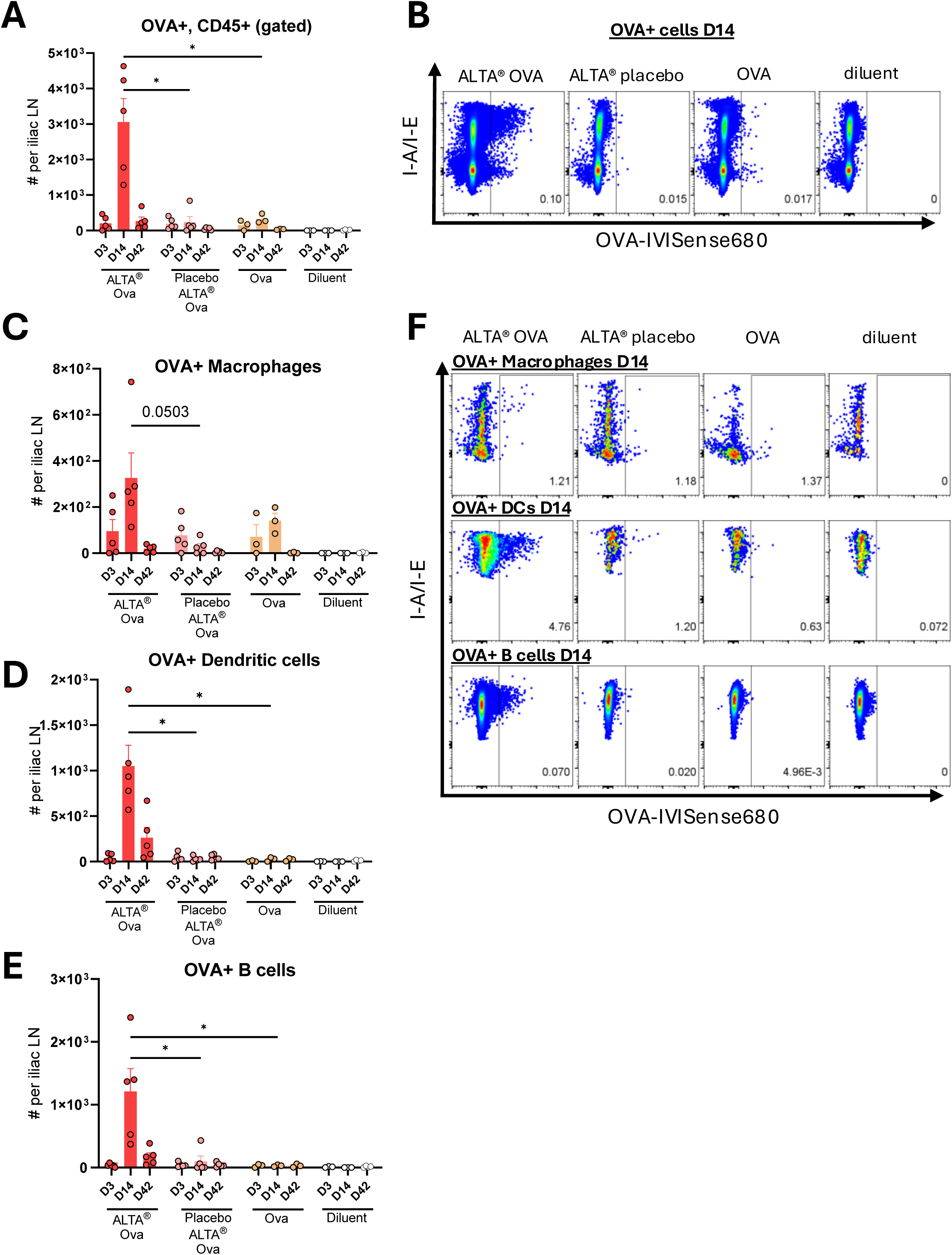
Antigen containment within ALTA^®^ particles results in an increased number of antigen-positive cells in the dLN. C57BL6 mice were i.m. immunized with OVA-IVISense680 dye conjugate formulated using ALTA^®^ platform (ALTA^®^ OVA) or OVA-IVISense680 dye conjugate mixed in diluent with placebo ALTA^®^ particles formulated without antigen (Placebo ALTA^®^). Liquid control groups were injected with liquid OVA-IVISense680 or diluent. All groups except diluent control were administered 820 ng OVA dose. At days 3, 7, and 42, iliac LN was processed and analyzed by flow cytometry. (**A**) Cell number of OVA-IVIS680+ CD45+ immune cells in iliac LN. Cell populations were gated on singlets, live cells, CD45+, OVA+. Absolute count of OVA+ CD45+ per iliac lymph nodes were determined as follows: [(%OVA+/100)*(%CD45+/ 100)*(live cell count)]. (**B**) Representative flow cytometry plots for percent of OVA+ of CD45+ cells in iliac LN at day 14 post treatments. (**C**–**E**) Absolute cell number of OVA+ APC subset per iliac LN. Cell number of OVA+ APCs iliac LN was determined as follows: [(%OVA+/100)*(%APC of CD45+/100)*(%CD45+/100)*(live cell count). Antigen-presenting cell populations were gated on singlets, live cells, CD45+, CD3- subsets ahead of the shown plots. (**C**) Macrophages were identified as CD45+, CD11b+, Ly6G-, Ly6c int/-, and F4/80+. (**D**) DCs were identified as CD45+, I-A/I-E+, CD11c^hi^. (**E**) B cells were identified as CD45+, I-A/I-E+, B220+. (**F**) Representative flow cytometry plots displaying percent of OVA+ of macrophages, DCs, B cells in iliac LN at day 14 post treatments. Shown are representative data from one of two experiments. N=3-5 /group. Mean ± SEM. Unpaired t test with Welch’s correction (shown are comparisons between select groups (ALTA® OVA vs ALTA® Placebo; ALTA® OVA vs OVA) at matching time points). *p ≤ 0.05.

When comparing ALTA^®^ OVA and ALTA^®^ placebo groups, the most apparent differences in the numbers of the total and antigen-positive cell subsets were observed in the dLN as compared to the site of injection. The increased presence of the total and antigen positive cell subsets in the dLN post administration demonstrates that containment of the antigen within particles facilitated the trafficking and/or maintenance of the antigen in the dLNs promoting antigen-specific adaptive immune responses.

### 3.9. Immunogenicity of ALTA^®^ vaccine containing clinically relevant vaccine antigens from *B. pseudomallei* and the adjuvant CpG ODN 2006

Next, the ALTA^®^ platform was applied to vaccine antigens derived from a facultative intracellular Gram-negative bacterium *B. pseudomallei*, the causative agent of melioidosis^29^. A combination of CPS-CRM197, Hcp1, and CpG ODN 2006 were formulated in proprietary buffer, spray-dried, and coated with 50 cycles of ALD to generate ALTA^®^ particles. To assess immunogenicity, C57BL6 mice were immunized with a range of ALTA^®^ doses (Supplementary Figure 7A). In this study, control mice were immunized with liquid formulations designed to match the antigen and adjuvant contents in ALTA^®^. Therefore, the liquid formulation contained equal amounts of antigens and CpG. Additionally, Alhydrogel^®^ was added in the control liquid formulation at a dose that matched the aluminum content of ALTA^®^ to account for any adjuvating effect of aluminum. Sera were collected at 4 weeks post immunizations to measure the anti-CPS and anti-Hcp1 IgG titers by ELISA. While mice immunized with the control liquid vaccine did not develop any detectable anti-CPS IgG response at low antigen doses (0.028 and 0.083 µg CPS), ALTA^®^ immunized mice elicited anti-CPS IgG at all doses tested (**Figure 9A**). These data show ALTA^®^ platform is suitable for use with a glycoconjugate antigen and that the immunogenicity of the antigen is improved when formulated using the ALTA^®^ platform. Previous work on *B. pseudomallei* vaccines showed IFN-γ secreting T cell responses to Hcp1 correlated with an improved survival of acute melioidosis patients^50^. Here, the anti-Hcp1 IgG titers were similar between the groups immunized with ALTA^®^ and the control liquid vaccine (**Figure 9B**). To measure the Hcp1-specific IFN-γ secreting T cell responses, splenocytes were stimulated with an Hcp1 peptide pool and analyzed by ELISpot. Starting at 0.5 µg Hcp1/dose and above, the number of IFN-γ-producing cells was higher in the ALTA^®^ immunized groups compared to the dose matched liquid control groups (**Figure 9C**). These data provide an example of the applicability of ALTA^®^ platform using disease-specific vaccine antigens (both protein and glycoconjugate) from *B. pseudomallei*. Next the immunogenicity of ALTA^®^ was compared against a liquid vaccine formulation containing higher adjuvant doses shown to provide protection in mice^31^. In addition, the same liquid control formulation described above containing matching antigen and adjuvant doses was used (Supplementary Figure 7B). All groups were immunized twice with equivalent antigen doses in a prime and boost scheduled 4 weeks apart. Sera were collected 2 weeks following the boost to measure anti-CPS and anti-Hcp1 IgG titers. All groups elicited robust anti-CPS IgG antibody responses (**Figure 9D**, E). The anti-Hcp1 IgG titers were equivalent between the groups immunized with ALTA^®^ or the liquid control containing higher adjuvant doses but were significantly reduced for the group immunized with the liquid formulation containing adjuvant doses matched to ALTA^®^ (**Figure 9F**, G). The T cell response as measured by IFN-γ producing splenocytes following Hcp1 peptide stimulation was highest in the group immunized with ALTA^®^ (**Figure 9H**). Thus, despite lower amounts of CpG or aluminum, the ALTA^®^ formulation generated robust humoral and cellular immune responses that were either similar or higher to a strongly adjuvanted liquid control. Taken together, these data demonstrate that when the ALTA^®^ platform was applied to the clinically relevant vaccine antigens, CPS-CRM197 and Hcp1, it can be adjuvant-sparing and elicit enhanced humoral and cell-mediated immunity.

**Figure 9.**
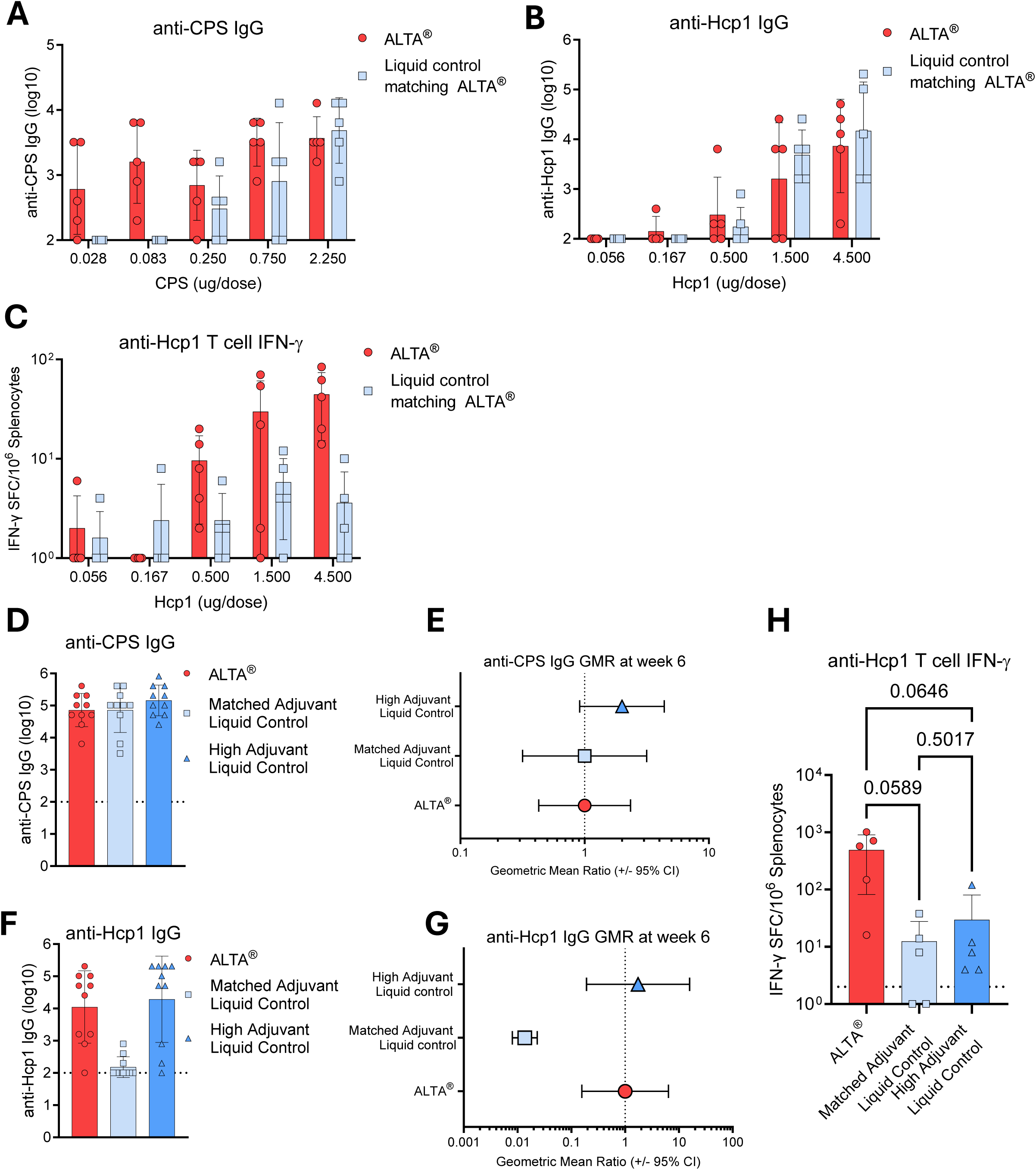
Immunogenicity of ALTA^®^ containing clinically relevant vaccine antigens from *B. pseudomallei* and the adjuvant CpG ODN 2006. A combination of antigens derived from *B. pseudomallei (*CPS and Hcp1) and CpG ODN 2006 were spray-dried and coated with 50 cycles of ALD to generate ALTA^®^ particles. (**A**-**C**) C57BL6 mice were s.c. immunized with a range of antigen doses formulated using ALTA^®^ platform. Control mice were immunized with liquid formulations designed to match the antigen and adjuvant contents in ALTA^®^, where Alhydrogel^®^ was used to match aluminum contents between conditions. (**A**–**C**) Sera were collected at 4 weeks post injections to measure the anti-CPS and anti-Hcp1 IgG titers by ELISA. The number of IFN-γ-producing cells was measured by ELISpot in spleens at week 4. (**A**) Anti-CPS IgG titer (log10). (**B**) Anti-Hcp1 IgG titer (log10). (**C**) Number of IFN-γ-producing splenocytes after *ex vivo* stimulation with Hcp1-derived peptides. (**D**–**H**) ALTA^®^ platform formulated antigens and CpG ODN 2006 or liquid controls were administered s.c. on day 0 and boosted on day 28. Antigen doses were matched between all groups. Adjuvant doses were: 10 ug CpG ODN 2006 + 250 ug Alhydrogel^®^ per dose of liquid control (High Adjuvant Liquid Control) and 0.38 µg CpG ODN 2006 + 3.5 µg Alhydrogel^®^ per dose of liquid control (Matched Adjuvant Liquid Control). Sera were collected at 6 weeks post prime to measure the anti-CPS and anti-Hcp1 IgG titers by ELISA. The number of IFN-γ-producing cells was measured by ELISpot in spleens at week 6. (**D, E**) Anti-CPS IgG titer (log10) (**D**) and GMR (**E**). (**F**, **G**) Anti-Hcp IgG1 titer (log10) (**F**) and GMR (**G**). (**H**) Number of IFN-γ-producing splenocytes after *ex vivo* stimulation with Hcp1-derived peptides. N=5-10 /group. Mean ± SD (**A**, **B**, **C**, **D**, **F**, **H**) or Geometric Mean Ratio ± 95% CI (**E**, **G**). Unpaired t test with Welch’s correction (p values shown) (**H**).

## 4. Discussion

### 4.1 Improved immunogenicity compared to clinically licensed vaccine adjuvants and their mimetics

Here, using both the model antigen OVA and clinically relevant *Burkholderia* antigens, the immunogenicity of the novel ALTA^®^ vaccine platform was characterized in comparison to liquid vaccine formulations adjuvanted with clinically licensed adjuvants or their mimetics. Vaccination with ALTA products resulted in a strong and durable humoral response and strikingly resulted in a more robust CD8+ immune response compared to the liquid vaccine formulations tested.

The humoral responses generated by the ALTA^®^ platform outperformed the liquid vaccine adjuvants and were indicative of a more balanced immune response. At low antigen doses, higher antigen-specific IgG1 titers were observed after immunization with ALTA^®^ than the adjuvanted liquid vaccine formulations, demonstrating an antigen-sparing property of the ALTA^®^ platform. With an increase in the antigen dose, the differences between the groups became less apparent, indicating a saturating response. ALTA^®^ immunized mice achieved high antibody production at significantly lower doses of aluminum than the Alhydrogel^®^ adjuvanted vaccine immunized mice. At all antigen doses tested, ALTA^®^ elicited higher IgG2c titers than those elicited by mice immunized with Alhydrogel^®^ or AddaVax™ adjuvanted liquid vaccine formulations. Furthermore, the ratio of IgG2c to IgG1 was calculated to characterize the Th1 versus Th2 skewing of the immune response post ALTA^®^ immunization^36^. Among all groups, vaccination with ALTA^®^ resulted in the production of both IgG1 and IgG2c at a ratio closest to one. These data demonstrated a more balanced Th1/Th2 response post administration of ALTA^®^ formulated antigen suggesting a possible broad applicability of the platform to vaccines against a wide range of pathogens.

The most striking difference in the immune responses to ALTA^®^ compared to other adjuvant formulations was the magnitude of a CD8+ T cell response. At all antigen doses and regiments tested, the magnitude of the CD8+ T cell response was higher after immunization with OVA formulated using ALTA^®^ platform than the OVA formulated with classical adjuvants. Here, ALTA^®^ surpassed vaccine formulations known to elicit cell-mediated responses (AddaVax™ or Ahydrogel^®^ + CpG ODN 1018) in the magnitude of the OVA-specific CD8+ T cell response^24,51^. Following one or two administrations of ALTA^®^ OVA, the OVA-specific CD8+ T cells were present in blood and spleens, demonstrated cytotoxic capacity (Granzyme B+), and produced cytokines upon antigen re-stimulation. Although less prominent than CD8+ T cells, the OVA-specific CD4+ T cells were also detected in the ALTA^®^ immunized mice. Based on these positive results using OVA as a model antigen, the use of the ALTA^®^ platform with antigens derived from an intracellular pathogen was further investigated.

Currently there are no licensed vaccines against meliodisis, but promising pre-clinical results are emerging from a liquid vaccine formulation given as 2-3 doses containing Alhydrogel^®^ and CpG^29^. Given these properties, it was hypothesized to be a good candidate for the ALTA^®^ platform. In this paper, *B. pseudomallei* antigens were formulated using the ALTA^®^ platform and improved immunogenicity was observed compared to liquid formulation controls. Specifically, immunization with ALTA^®^ containing the antigens CPS-CRM197 and Hcp1 elicited robust antibody and T cell IFN-γ production. Compared to dose-matched liquid controls, ALTA^®^ immunized mice developed anti-CPS IgG responses at lower doses of the glycoconjugate. Furthermore, ALTA^®^ immunized mice had higher numbers of anti-Hcp1 IFN-γ-producing cells than mice immunized with the liquid formulations. Overall, immunization with ALTA^®^ platform containing clinically relevant bacterial antigens elicited strong humoral and cell-mediated immunity in mice. Determining whether these findings will translate into larger animal species is of interest and those studies are in pursuit.

### 4.2. Mechanisms of Action: Key attributes of ALTA^®^ platform which may improve immune response

#### Extended antigen presence in vivo

The enhanced immunogenicity of the antigens formulated using the ALTA^®^ platform could be associated with the extended presence of the antigens *in vivo*. The release of the contained antigen from ALTA^®^ particles is dependent on the number of ALD cycles, and the timing of antibody response *in vivo* correlates with coat thickness^1,4,7^. Although ALTA^®^ particles used in this study were coated with a relatively low number of ALD cycles (50), longitudinal *in vivo* imaging of mice showed the fluorescent signal from the 50-cycle ALTA^®^ containing OVA-IVISense680 was detectable for several weeks (Strand et. al. manuscript in preparation). The flow cytometry analysis presented in the current study showed the fluorescent antigen post ALTA^®^ OVA-IVISense680 i.m. administration was detectable at the SOI and in the dLNs for the duration of the study (6 weeks). Furthermore, the numbers of antigen-positive cells were significantly higher two weeks after administration of fluorescent OVA formulated in ALTA^®^ platform than liquid OVA administered alone or with placebo ALTA^®^. Multiple studies showed that prolonged bioavailability of the antigen post vaccination coinciding with the germinal center (GC) response promoted humoral immunity^11,12,52–54^. For example, enhanced binding of immunogens to alum for prolonged bioavailability *in vivo* resulted in an increased humoral response compared to conventional alum-adsorbed adjuvants^55^. Other slow release vaccine approaches, such as repeated injections, two-dose “extended priming”^10^, miniosmotic pumps^12^, microneedle patches^11^, as well as innovative biomaterials, including hydrogels^56–59^, biodegradable polymer microparticles^60^, slow-release alum^61^ also resulted in enhanced humoral responses relative to traditional bolus immunization strategies^13,52^. In general, the slow release vaccines enhance the GC response^62^, including activation of the T follicular helper cells (Tfh), follicular DCs, and GC B cells. Thus, one of the mechanisms by which ALTA^®^ platform enhances antibody production could be associated with the extended antigen bioavailability and a more efficient GC response. Future studies will be directed onto investigating the GC response post ALTA^®^ vaccination in more detail.

#### Antigen containment within particles

Encapsulation of antigens into the spray-dried core of the ALTA^®^ particles may explain their improved immunogenicity over liquid formulations with adjuvants. In general, particulate antigens were shown to be more immunogenic than their soluble counterparts ^63–65^. Vaccine antigens can be delivered via particles either by adsorption to the surface or by encapsulation into the core of particles. Most antigen formulated using the ALTA^®^ platform is encapsulated (a small percent is immediately available likely due to imperfections in research scale production). While immunogenicity of the protein antigens attached to the surface of particles was described in detail^66^, examples of vaccine technologies utilizing microparticles with contained antigens are limited. A few studies showed encapsulation of vaccine antigens into particles improved humoral and cell-mediated immune responses^65,67,68^. For example, the vaccine consisting of β-glucan microparticles encapsulating antigen and the aluminum hydroxide colloid strongly increased DC activation, production of IgG1 and IgG2a, and IFN-γ+ CD8+ T cells, in addition to lowering tumor growth in mice^67^. Perhaps the most common approach for encapsulating vaccine antigens into microparticles is poly(lactic-co-glycolic acid) (PLGA)-based. PLGA microparticles were shown to increase antigen uptake and activation of APCs, antibody titers (including IgG2a) and T cell responses^68,69^. Overall, vaccines containing particles were shown to induce strong cell-mediated immunity associated with a more efficient uptake of the antigens by the phagocytes and enhanced antigen presentation, including cross-presentation.

The current study demonstrates that encapsulation of antigens is a critical attribute of the ALTA^®^ platform and essential for its immunogenicity. Compared to antigen formulated within spray-dried core of the ALTA^®^ platform, administration of the antigen mixed with placebo ALTA^®^ particles elicited significantly lower antibody and T cell responses. In addition, the numbers of the antigen-positive cells in tissues, including injected muscle and dLN, were also reduced in that group. Interestingly, higher numbers of antigen-positive cells were found at the site of injection following administration of OVA mixed with placebo ALTA^®^ compared to administration of OVA alone. This result demonstrated that ALTA^®^ particles may have an intrinsic adjuvating capacity. However, only ALTA^®^ containing encapsulated antigen led to a prolonged presence of the antigen-positive cells at the site of injection and their significant increase in the dLN. These data suggest that when the antigen is contained within the alumina coating, the ALTA^®^ platform may extend delivery and increase uptake of the antigen to APCs from the site of injection and facilitate antigen re-localization to the dLNs. Therefore, while ALTA^®^ particles demonstrate an intrinsic adjuvating capacity, ALTA^®^ platform containing encapsulated antigen may act as an antigen delivery system.

In summary, the ALTA^®^ platform is unique because it combines features of several vaccine technologies. Content-wise, ALTA^®^ particles contain aluminum, a shared feature with other adjuvants in clinical use. This feature may contribute to an intrinsic adjuvating capacity of ALTA^®^ particles as well as an enhancement of the humoral response. Second, the ALTA^®^ platform allows a temporal control of antigen release and extends bioavailability of antigens *in vivo* - an important feature shared with other controlled release vaccine technologies. This feature may contribute to an overall improved immunogenicity of the platform, especially enhancement of the humoral immunity. Third, ALTA^®^ technology allows for encapsulation of antigens/adjuvants in the spray dried core of microparticles, which may be beneficial for both humoral and cellular immune responses. Continued efforts to further understand the mechanisms underlaying ALTA^®^ platform immunogenicity will aid in the clinical development of a more effective and safer vaccine.

## 4. Conclusion

ALTA^®^ vaccine technology combines spray drying and ALD to create a thermostable product with unique features that facilitate an adaptive immune response. These studies demonstrate that ALTA^®^ generates both humoral and cellular immune responses that are comparable to or offer improvement over the responses generated by formulations containing a set of commonly used liquid adjuvants. Although further investigation is still needed, the balanced and durable immune responses that were observed in these studies using lower doses of antigen and/or adjuvant demonstrate the potential for this technology to have a significant impact on public health

## Author contributions

D. L. I. and S. W. B. conceptualized studies. D. L. I., M. S. L, A. B. C., I. R. W. conducted experiments, generated data and interpreted experimental results. E. M. S., K. A. S. conducted experiments and generated data. L. R. A., S. S., S. B. W., F. U., M. N. B., P. J. B. provided reagents, conducted experiments and generated data related to *B. pseudomalei*. D. L. I. prepared the original draft manuscript. D. L. I., S. W. B., K. A. S., M. S. L., M. N. B., P. J. B. provided review and editing.

## Acknowledgements

The authors would like to acknowledge and thank the following contributors: Antu K. Dey for reviewing the manuscript. Kimberly Erickson for reviewing the manuscript and supervision of the studies related to *B. pseudomallei.* Emma Palm, Grace Beck, Brittany Villanueva, Edis Cehic, and Sarah Adzema for contributions with manufacturing the ALTA^®^ products used in these studies. Emily Hite for conjugation of OVA and IVISense680 probe. Jason Yau and Cisloynny Beauchamp-Perez for analytical work on ALTA^®^ formulated vaccine against *B. pseudomallei* and preparation of liquid controls. Lindsay Larson (Office of Animal Resources- CU Boulder), Heather D’Angelo and Ashley Gerwing for assistance with murine injections and bleeds. Giovanny Hernandez for assistance with tissue harvests to evaluate T cell responses. Jacob Loughry (UNR) for assistance with antigen production, mouse studies, and immune assays.

The research described in this report was funded by both private equity and from grants distributed by the Gates Foundation (INV-064824 and INV-076862) and the Defense Treat Reduction Agency (HDTRA1-23-C-0009). S.B.W, F.U.M, S.S., M.N.B and P.J.B were supported in part by HDTRA1-23-C-0009 (to VitriVax) HDTRA1-18-C-0062 (to UNR). S.S. was supported in part by the Springboard Science and Research Fund, University of Nevada, Reno School of Medicine. Funders played no role in study design, data collection, analysis and interpretation of data, or the writing of this manuscript.

## Data availability

The datasets generated and/or analyzed during the current study are not publicly available due to company policy but will be made available from the corresponding author on request.

## References

1. Brubaker, S. W. et al. Demonstration of Tunable Control over a Delayed-Release Vaccine Using Atomic Layer Deposition. Vaccines 12, 761 (2024).

2. Hassett, K. J. et al. Glassy-state stabilization of a dominant negative inhibitor anthrax vaccine containing aluminum hydroxide and glycopyranoside lipid A adjuvants. J. Pharm. Sci. 104, 627–639 (2015).

3. Clausi, A. L., Merkley, S. A., Carpenter, J. F. & Randolph, T. W. Inhibition of aggregation of aluminum hydroxide adjuvant during freezing and drying. J. Pharm. Sci. 97, 2049–2061 (2008).

4. Garcea, R. L. et al. Single-administration, thermostable human papillomavirus vaccines prepared with atomic layer deposition technology. NPJ Vaccines 5, 45 (2020).

5. Weimer, A. W. Particle atomic layer deposition. J. Nanoparticle Res. Interdiscip. Forum Nanoscale Sci. Technol. 21, 9 (2019).

6. Randolph, T. et al. Superior immune responses from thermostable, single-administration rabies vaccines prepared using atomic layer deposition. J. Pharm. Sci. 114, 103936 (2025).

7. Cohen, A. A. et al. Broad anti-sarbecovirus responses elicited by a single administration of mosaic-8 RBD-nanoparticle vaccine prepared using atomic layer deposition. iScience 28, 113649 (2025).

8. Coleman, H. J. et al. Lipid-free, thermostable mRNA vaccines prepared using atomic layer deposition. J. Pharm. Sci. 104066 (2025) doi:10.1016/j.xphs.2025.104066.

9. Duralliu, A. et al. The influence of the closure format on the storage stability and moisture content of freeze-dried influenza antigen. Vaccine 37, 4485–4490 (2019).

10. Bhagchandani, S. H. et al. Two-dose ‘extended priming’ immunization amplifies humoral immune responses by synchronizing vaccine delivery with the germinal center response. BioRxiv Prepr. Serv. Biol. 2023.11.20.563479 (2023) doi:10.1101/2023.11.20.563479.

11. Boopathy, A. V. et al. Enhancing humoral immunity via sustained-release implantable microneedle patch vaccination. Proc. Natl. Acad. Sci. U. S. A. 116, 16473–16478 (2019).

12. Cirelli, K. M. et al. Slow Delivery Immunization Enhances HIV Neutralizing Antibody and Germinal Center Responses via Modulation of Immunodominance. Cell 177, 1153–1171.e28 (2019).

13. Roth, G. A. et al. Designing spatial and temporal control of vaccine responses. Nat. Rev. Mater. 7, 174–195 (2022).

14. Lan, J., Feng, D., He, X., Zhang, Q. & Zhang, R. Basic Properties and Development Status of Aluminum Adjuvants Used for Vaccines. Vaccines 12, 1187 (2024).

15. Laera, D., HogenEsch, H. & O’Hagan, D. T. Aluminum Adjuvants-’Back to the Future’. Pharmaceutics 15, 1884 (2023).

16. Aung, A. & Irvine, D. J. Modulating Antigen Availability in Lymphoid Organs to Shape the Humoral Immune Response to Vaccines. J. Immunol. Baltim. Md 1950 212, 171–178 (2024).

17. Chen, K. et al. Harnessing cellular immunity for next-generation vaccines against respiratory viruses: mechanisms, platforms, and optimization strategies. Front. Immunol. 16, 1618406 (2025).

18. Shapiro, J. R., Corrado, M., Perry, J., Watts, T. H. & Bolotin, S. The contributions of T cell-mediated immunity to protection from vaccine-preventable diseases: A primer. Hum. Vaccines Immunother. 20, 2395679 (2024).

19. Borgo, G. M. & Rutishauser, R. L. Generating and measuring effective vaccine-elicited HIV-specific CD8 + T cell responses. Curr. Opin. HIV AIDS 18, 331–341 (2023).

20. Nogimori, T. et al. Functional changes in cytotoxic CD8+ T-cell cross-reactivity against the SARS-CoV-2 Omicron variant after mRNA vaccination. Front. Immunol. 13, 1081047 (2022).

21. Peng, S. et al. Particulate Alum via Pickering Emulsion for an Enhanced COVID-19 Vaccine Adjuvant. Adv. Mater. Deerfield Beach Fla 32, e2004210 (2020).

22. O’Hagan, D. T., Ott, G. S., De Gregorio, E. & Seubert, A. The mechanism of action of MF59 - an innately attractive adjuvant formulation. Vaccine 30, 4341–4348 (2012).

23. Cantisani, R. et al. Vaccine adjuvant MF59 promotes retention of unprocessed antigen in lymph node macrophage compartments and follicular dendritic cells. J. Immunol. Baltim. Md 1950 194, 1717–1725 (2015).

24. Kim, E. H. et al. Squalene emulsion-based vaccine adjuvants stimulate CD8 T cell, but not antibody responses, through a RIPK3-dependent pathway. eLife 9, e52687 (2020).

25. Dong, M., Meinerz, N. M., Walker, K. D., Garcea, R. L. & Randolph, T. W. Thermostability of a trivalent, capsomere-based vaccine for human papillomavirus infection. Eur. J. Pharm. Biopharm. OM. J. Arbeitsgemeinschaft Pharm. Verfahrenstechnik EV 168, 131–138 (2021).

26. Witeof, A. E. et al. Atomic-Layer Deposition Processes Applied to Phage λ and a Phage-like Particle Platform Yield Thermostable, Single-Shot Vaccines. J. Pharm. Sci. 111, 1354–1362 (2022).

27. Witeof, A. E. et al. A Single Dose, Thermostable, Trivalent Human Papillomavirus Vaccine Formulated Using Atomic Layer Deposition. J. Pharm. Sci. 112, 2223–2229 (2023).

28. Coleman, H. J. et al. Formulation of three tailed bacteriophages by spray-drying and atomic layer deposition for thermal stability and controlled release. J. Pharm. Sci. 113, 3238–3245 (2024).

29. Sengyee, S., Weiby, S. B., Rok, I. T., Burtnick, M. N. & Brett, P. J. Melioidosis vaccines: recent advances and future directions. Front. Immunol. 16, 1582113 (2025).

30. Burtnick, M. N. et al. Development of Subunit Vaccines That Provide High-Level Protection and Sterilizing Immunity against Acute Inhalational Melioidosis. Infect. Immun. 86, e00724–17 (2018).

31. Biryukov, S. S. et al. Comparison of homologous and heterologous vaccination strategies for combating disease caused by Burkholderia pseudomallei. Front. Immunol. 16, 1596265 (2025).

32. Schmidt, L. K. et al. Development of Melioidosis Subunit Vaccines Using an Enzymatically Inactive Burkholderia pseudomallei AhpC. Infect. Immun. 90, e0022222 (2022).

33. Biryukov, S. S. et al. Evaluation of two different vaccine platforms for immunization against melioidosis and glanders. Front. Microbiol. 13, 965518 (2022).

34. Arunachalam, P. S. et al. Durability of immune responses to mRNA booster vaccination against COVID-19. J. Clin. Invest. 133, e167955 (2023).

35. Tzeng, T.-T. et al. A TLR9 agonist synergistically enhances protective immunity induced by an Alum-adjuvanted H7N9 inactivated whole-virion vaccine. Emerg. Microbes Infect. 12, 2249130 (2023).

36. Nazeri, S., Zakeri, S., Mehrizi, A. A., Sardari, S. & Djadid, N. D. Measuring of IgG2c isotype instead of IgG2a in immunized C57BL/6 mice with Plasmodium vivax TRAP as a subunit vaccine candidate in order to correct interpretation of Th1 versus Th2 immune response. Exp. Parasitol. 216, 107944 (2020).

37. Cribbs, D. H. et al. Adjuvant-dependent modulation of Th1 and Th2 responses to immunization with beta-amyloid. Int. Immunol. 15, 505–514 (2003).

38. Grigoryan, L. et al. Adjuvanting a subunit SARS-CoV-2 vaccine with clinically relevant adjuvants induces durable protection in mice. NPJ Vaccines 7, 55 (2022).

39. Fong, Y. et al. Immune correlates analysis of the PREVENT-19 COVID-19 vaccine efficacy clinical trial. Nat. Commun. 14, 331 (2023).

40. Violán, C. et al. Immune Durability and Breakthrough Infections 15 Months After SARS-CoV-2 Boosters in People over 65: The IMMERSION Study. Vaccines 13, 738 (2025).

41. da Silva Antunes, R., et al. Evolution of SARS-CoV-2 T cell responses as a function of multiple COVID-19 boosters. Cell Rep. 44, 115907 (2025).

42. Seder, R. A. & Hill, A. V. Vaccines against intracellular infections requiring cellular immunity. Nature 406, 793–798 (2000).

43. Kalimuddin, S. et al. Vaccine-induced T cell responses control Orthoflavivirus challenge infection without neutralizing antibodies in humans. Nat. Microbiol. 10, 374–387 (2025).

44. Bange, E. M. et al. CD8+ T cells contribute to survival in patients with COVID-19 and hematologic cancer. Nat. Med. 27, 1280–1289 (2021).

45. Peng, K., Zhao, X., Fu, Y.-X. & Liang, Y. Eliciting antitumor immunity via therapeutic cancer vaccines. Cell. Mol. Immunol. 22, 840–868 (2025).

46. Kaech, S. M. & Cui, W. Transcriptional control of effector and memory CD8+ T cell differentiation. Nat. Rev. Immunol. 12, 749–761 (2012).

47. Mahajan, S. D., Aalinkeel, R., Schwartz, S. A., Chawda, R. P. & Nair, M. P. N. Effector cell mediated cytotoxicity measured by intracellular Granzyme B release in HIV infected subjects. Biol. Proced. Online 5, 182–188 (2003).

48. Harrell, M. I., Iritani, B. M. & Ruddell, A. Lymph node mapping in the mouse. J. Immunol. Methods 332, 170–174 (2008).

49. Ding, Y., Li, Z., Jaklenec, A. & Hu, Q. Vaccine delivery systems toward lymph nodes. Adv. Drug Deliv. Rev. 179, 113914 (2021).

50. Sengyee, S. et al. Melioidosis Patient Survival Correlates With Strong IFN-γ Secreting T Cell Responses Against Hcp1 and TssM. Front. Immunol. 12, 698303 (2021).

51. Li, Y. & Chen, X. CpG 1018 Is an Effective Adjuvant for Influenza Nucleoprotein. Vaccines 11, 649 (2023).

52. Tam, H. H. et al. Sustained antigen availability during germinal center initiation enhances antibody responses to vaccination. Proc. Natl. Acad. Sci. U. S. A. 113, E6639–E6648 (2016).

53. Cirelli, K. M. & Crotty, S. Germinal center enhancement by extended antigen availability. Curr. Opin. Immunol. 47, 64–69 (2017).

54. Rodrigues, K. A. et al. Optimization of an alum-anchored clinical HIV vaccine candidate. NPJ Vaccines 8, 117 (2023).

55. Moyer, T. J. et al. Engineered immunogen binding to alum adjuvant enhances humoral immunity. Nat. Med. 26, 430–440 (2020).

56. Roth, G. A. et al. Injectable Hydrogels for Sustained Codelivery of Subunit Vaccines Enhance Humoral Immunity. ACS Cent. Sci. 6, 1800–1812 (2020).

57. Roth, G. A. et al. Prolonged Codelivery of Hemagglutinin and a TLR7/8 Agonist in a Supramolecular Polymer-Nanoparticle Hydrogel Enhances Potency and Breadth of Influenza Vaccination. ACS Biomater. Sci. Eng. 7, 1889–1899 (2021).

58. Gale, E. C. et al. Hydrogel-Based Slow Release of a Receptor-Binding Domain Subunit Vaccine Elicits Neutralizing Antibody Responses Against SARS-CoV-2. Adv. Mater. Deerfield Beach Fla 33, e2104362 (2021).

59. Hao, H. et al. Immunization against Zika by entrapping live virus in a subcutaneous self-adjuvanting hydrogel. *Nat*. Biomed. Eng. 7, 928–942 (2023).

60. Zhang, Y. et al. Evaluation of HPV-loaded PLGA microparticles as single-dose HPV vaccine: Insights for sustained-release vaccine development. Vaccine 55, 127024 (2025).

61. Rodrigues, K. A. et al. Vaccines combining slow release and follicle targeting of antigens increase germinal center B cell diversity and clonal expansion. Sci. Transl. Med. 17, eadw7499 (2025).

62. Ray, S., Puente, A., Steinmetz, N. F. & Pokorski, J. K. Recent advancements in single dose slow-release devices for prophylactic vaccines. Wiley Interdiscip. Rev. Nanomed. Nanobiotechnol. 15, e1832 (2023).

63. Scheicher, C., Mehlig, M., Dienes, H. P. & Reske, K. Uptake of bead-adsorbed versus soluble antigen by bone marrow derived dendritic cells triggers their activation and increases their antigen presentation capacity. Adv. Exp. Med. Biol. 378, 253–255 (1995).

64. Scheicher, C., Mehlig, M., Dienes, H. P. & Reske, K. Uptake of microparticle-adsorbed protein antigen by bone marrow-derived dendritic cells results in up-regulation of interleukin-1 alpha and interleukin-12 p40/p35 and triggers prolonged, efficient antigen presentation. Eur. J. Immunol. 25, 1566–1572 (1995).

65. Koppolu, B. & Zaharoff, D. A. The effect of antigen encapsulation in chitosan particles on uptake, activation and presentation by antigen presenting cells. Biomaterials 34, 2359–2369 (2013).

66. Van Braeckel-Budimir, N., Haijema, B. J. & Leenhouts, K. Bacterium-like particles for efficient immune stimulation of existing vaccines and new subunit vaccines in mucosal applications. Front. Immunol. 4, 282 (2013).

67. Liu, H. et al. Aluminum hydroxide colloid vaccine encapsulated in yeast shells with enhanced humoral and cellular immune responses. Biomaterials 167, 32–43 (2018).

68. Joshi, V. B., Geary, S. M. & Salem, A. K. Biodegradable particles as vaccine antigen delivery systems for stimulating cellular immune responses. Hum. Vaccines Immunother. 9, 2584–2590 (2013).

69. Kasturi, S. P. et al. Programming the magnitude and persistence of antibody responses with innate immunity. Nature 470, 543–547 (2011).

