## Supplemental Material for "Subunit Vaccination Using Atomic Layering Thermostable Antigen and Adjuvant (ALTA^®^) Platform Elicits Enhanced Humoral and Cellular Immune Responses"

### Supplementary Figure 1

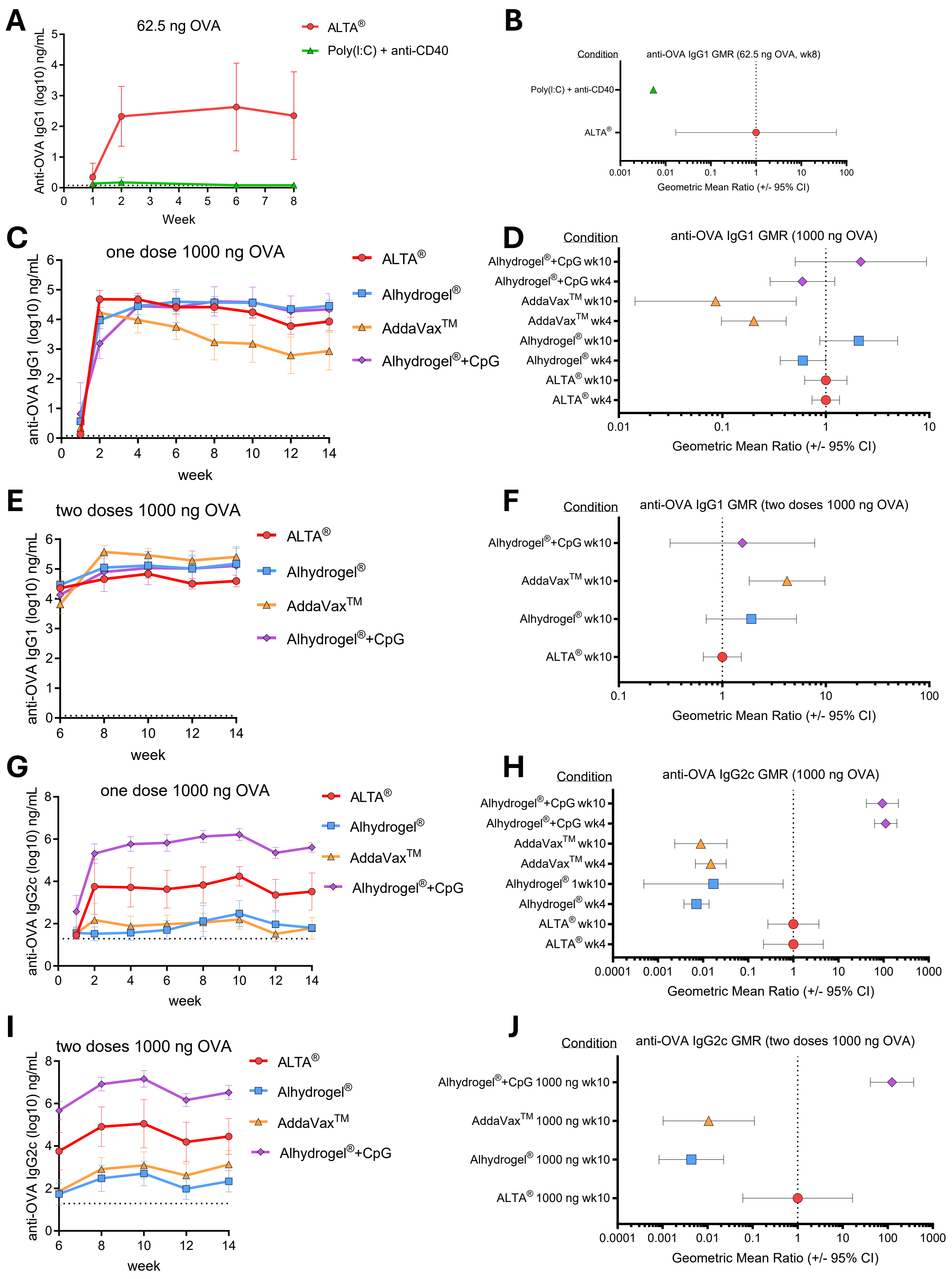

**Supplementary Figure 1. Administration of OVA antigen formulated using ALTA® platform elicits robust and Th1/Th2 balanced antibody production.**

(**A**, **B**) C57BL6 mice were injected i.m. with one dose of 62.5 ng of OVA antigen delivered in ALTA® platform or with poly(I:C)+anti-CD40. Shown are the anti-OVA IgG1 titers (log10) over 8 weeks (**A**) and GMR at week 8 (**B**). N=5 /group. (**C-J**) C57BL6 mice were injected i.m. with one (**C**, **D** and **G**, **H**) or two doses (**E**, **F** and **I**, **J**) of 1000 ng of OVA antigen delivered in ALTA® platform or with adjuvants, Alhydrogel®, AddaVax™, Alhydrogel®+CpG ODN 1018. (**C**) Anti-OVA IgG1 titers (log10) over 14 weeks and (**D**) the GMR at week 4 (n=9-10) and 10 (n=3-5) (1000 ng OVA). (**E**) Anti-OVA IgG1 titers (log10) before and after boost dose administered at week 6 and (**F**) the GMR at week 4 post boost (1000 ng OVA/dose). (**G**) Anti-OVA IgG2c titers (log10) over 14 weeks and (**H**) the GMR at week 4 (n=9-10) and 10 (n=3-5) (1000 ng OVA). (**I**) Anti-OVA IgG2c titers (log10) before and after boost dose administered at week 6 and (**J**) the GMR at week 4 post boost (n=4-5) (1000 ng OVA/dose). Mean ± SD (**A**, **C**, **E**, **G**, **I**) or Geometric Mean Ratio ± 95% CI (**B**, **D**, **F**, **H**, **J**).

### Supplementary Figure 2

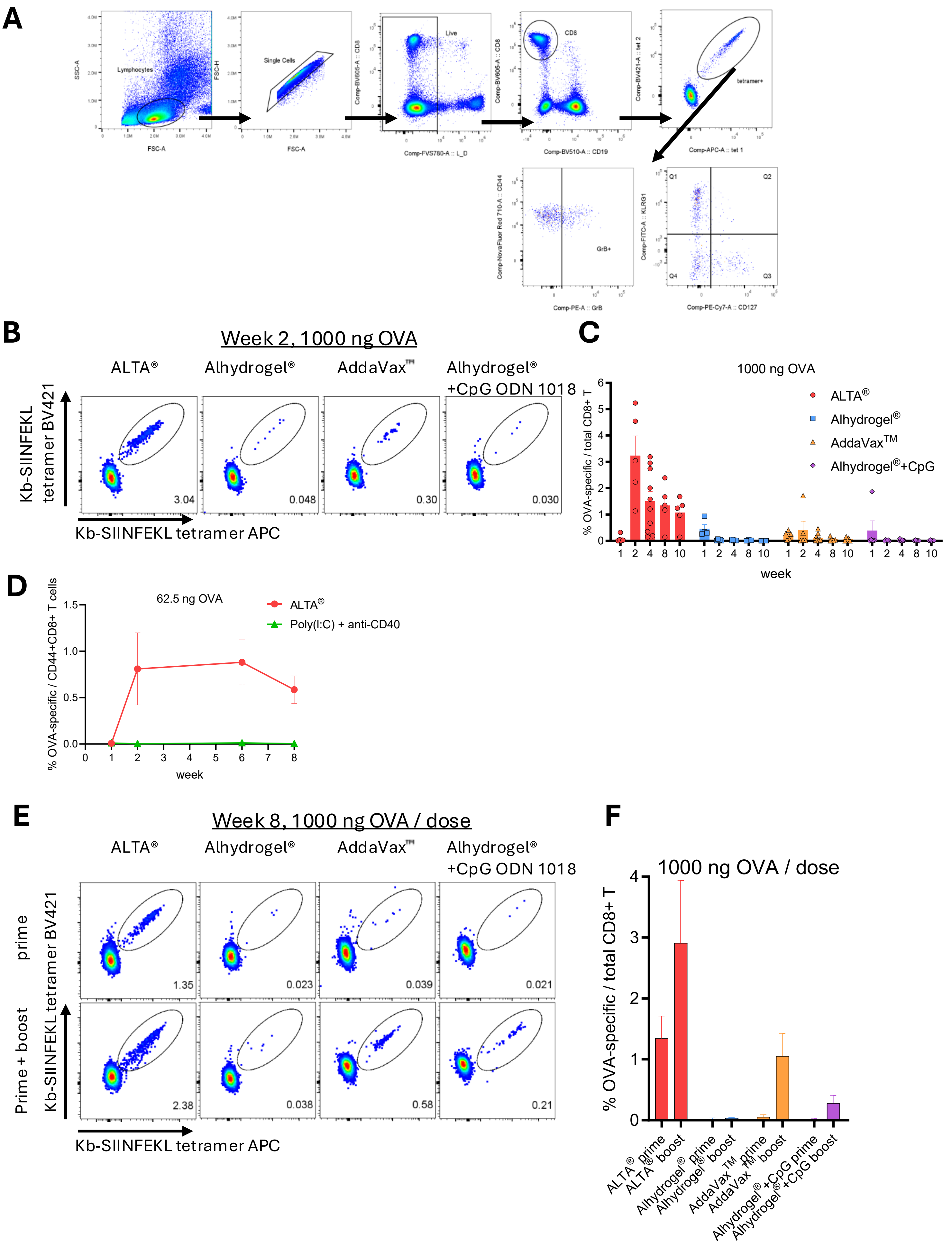

**Supplementary Figure 2. Administration of antigen formulated using ALTA® platform elicits a more robust antigen (OVA)-specific CD8+ T cell response than Alhydrogel®, AddaVax™, Alhydrogel®+CpG ODN 1018, poly(I:C)+anti-CD40.**

(**A**) Gating strategy for the identification and characterization of the OVA-specific CD8+ T cells in blood. The OVA-specific CD8+ T cells were gated as H-2K$^b$-SIINFEKL tetramer APC+ H-2K$^b$-SIINFEKL tetramer BV421+ in CD8α+CD19- live lymphocytes. (**B**, **C**) C57BL6 mice were i.m. immunized with one dose of 1000 ng of OVA antigen delivered in ALTA® or with adjuvants, including Alhydrogel®, AddaVax™, Alhydrogel®+CpG ODN 1018. Blood was collected for 10 weeks and analyzed for the presence of OVA-specific CD8+ T cells by flow cytometry. N=3-10 /group. (**B**) Representative dot plots depicting the percentage of OVA-specific CD8+ T cells of total CD8+ T cells at week 2 post-vaccination. (**C**) The frequency of OVA-specific CD8+ T cells in blood. (**D**) C57BL6 mice were i.m. immunized with 62.5 ng of OVA delivered in ALTA® or with poly(I:C)+anti-CD40. Blood was collected at weeks 1, 2, 6, 8 and analyzed for the percentage of OVA-specific CD8+ T cells (of CD44+ CD8+ T cells). N=5/group. (**E**, **F**) C57BL6 mice were i.m. immunized with one or two doses of 1000 ng of OVA delivered in ALTA® or with adjuvants, including Alhydrogel®, AddaVax™, Alhydrogel®+CpG ODN 1018. N=3-5/group. (**E**) Representative dot plots depicting the percentage of OVA-specific CD8+ T cells at week 8 post prime (top row) and 2 weeks after boost at week 6 (bottom row). (**F**) The frequency of OVA-specific CD8+ T cells at week 8 post prime and 2 weeks after week 6 boost with 1000 ng OVA. Mean ± SEM.

### Supplementary Figure 3

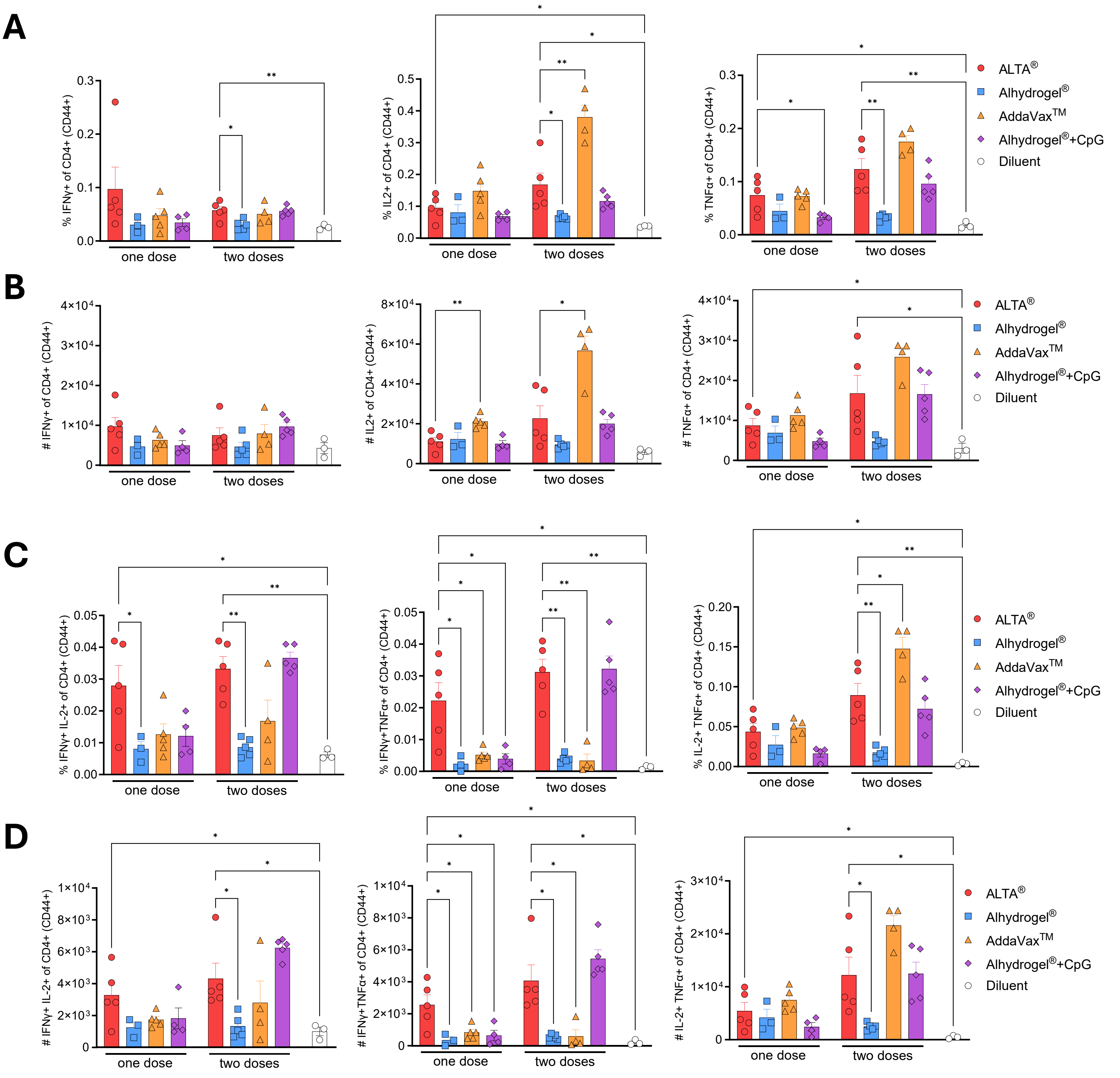

**Supplementary Figure 3. The cytokine-producing OVA-specific CD4+ T cells are present in the spleens of the ALTA® immunized mice.**

C57BL6 mice were i.m. immunized with 1000 ng of OVA antigen delivered in ALTA® or with Alhydrogel®, AddaVax™, Alhydrogel®+CpG ODN 1018. Indicated groups were boosted at week 6 post prime. The spleens were harvested at week 14 post prime and splenocytes were stimulated with the OVA peptide pool for 5 hours *ex vivo*. The expression of IFN-γ, IL-2, and/or TNF-α by CD4+ T cells (gated as CD44+CD4+CD19- live lymphocytes) was measured by intracellular cytokine staining. The frequency (**A**) and number per spleen (**B**) of the IFN-γ, IL-2, or TNF-α –expressing CD4+ T cells. (**C**, **D**) The frequency (**C**) and number per spleen (**D**) of the CD4+ T cells expressing two cytokines (IFN-γ+IL-2+, IFN-γ+TNF-α+, IL-2+TNF-α+). N=3-5/ group. Mean ± SEM. Unpaired t test with Welch's correction. *$p \leq 0.05$, **$p \leq 0.01$.

### Supplementary Figure 4

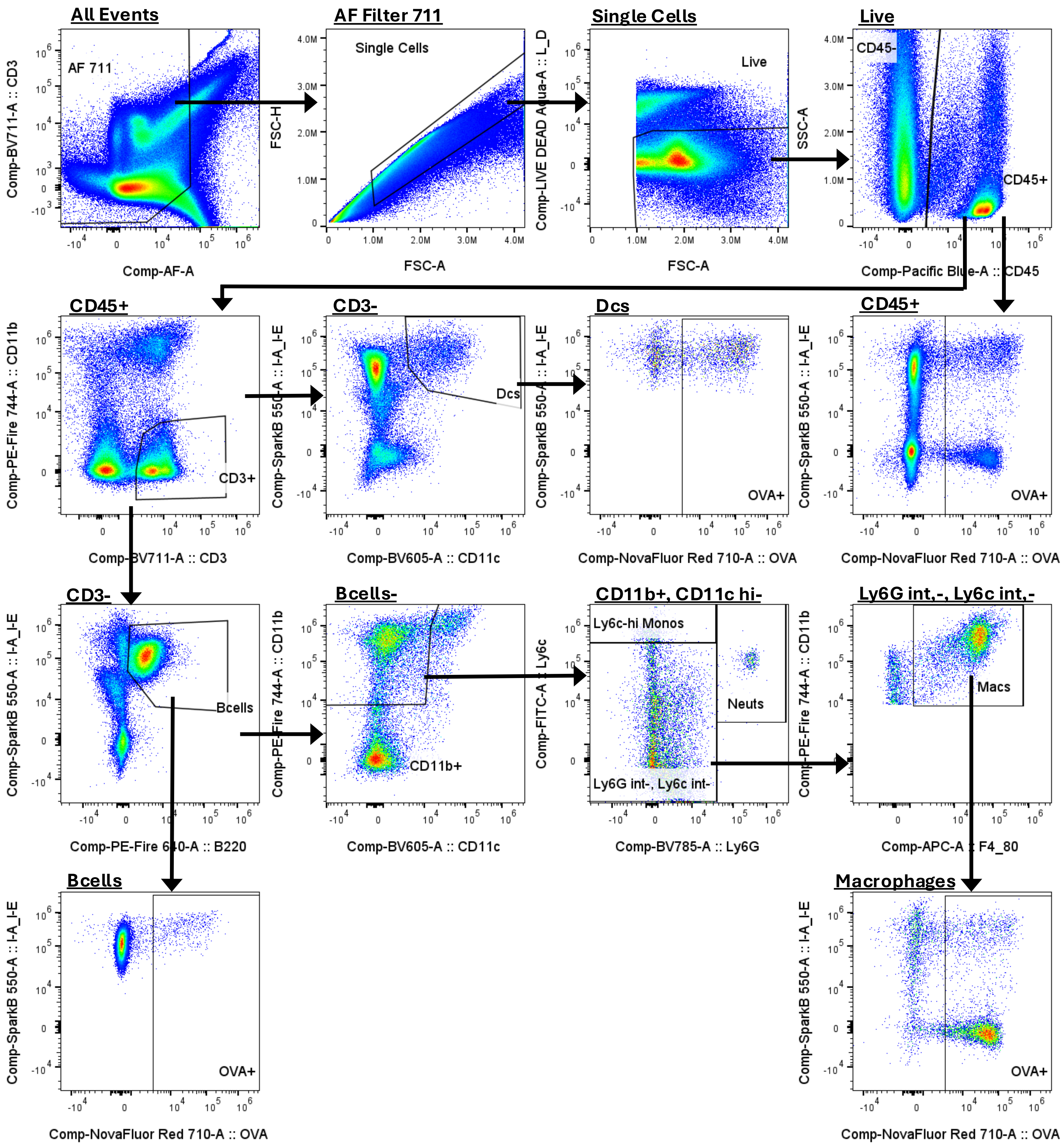

**Supplementary Figure 4. Gating strategy for phenotyping of the main immune cell subsets and identification of antigen-positive cells at the site of injection and dLN.**

For all analyses gating began with autofluorescence exclusion, then singlet and live cell selection. Next CD45+ and CD45- cell populations were gated. Of the CD45+ population, CD3+ T cells were identified and a T cell "Not" gate was created to remove the T cells from downstream analyses. B cells were then distinguished from the T cell population by I-A/I-E+ and B220+. Myeloid cells were then identified from the T cell and B cell negative population by CD11b+ and CD11c-, int. Neutrophils were identified as $Ly6G^{hi}$, Ly6c+ and inflammatory monocytes were identified as $Ly6c^{hi}$, Ly6G-, int. Macrophages were then distinguished from the Ly6c-, int, Ly6G-, int population as F4/80+. Dendritic cells were identified as I-A/I-E+, CD11c+ as a daughter population of T cells. All immune subsets were further analyzed for the presence of OVA-IVISense680. Analysis was performed in FlowJo software (BD).

### Supplementary Figure 5

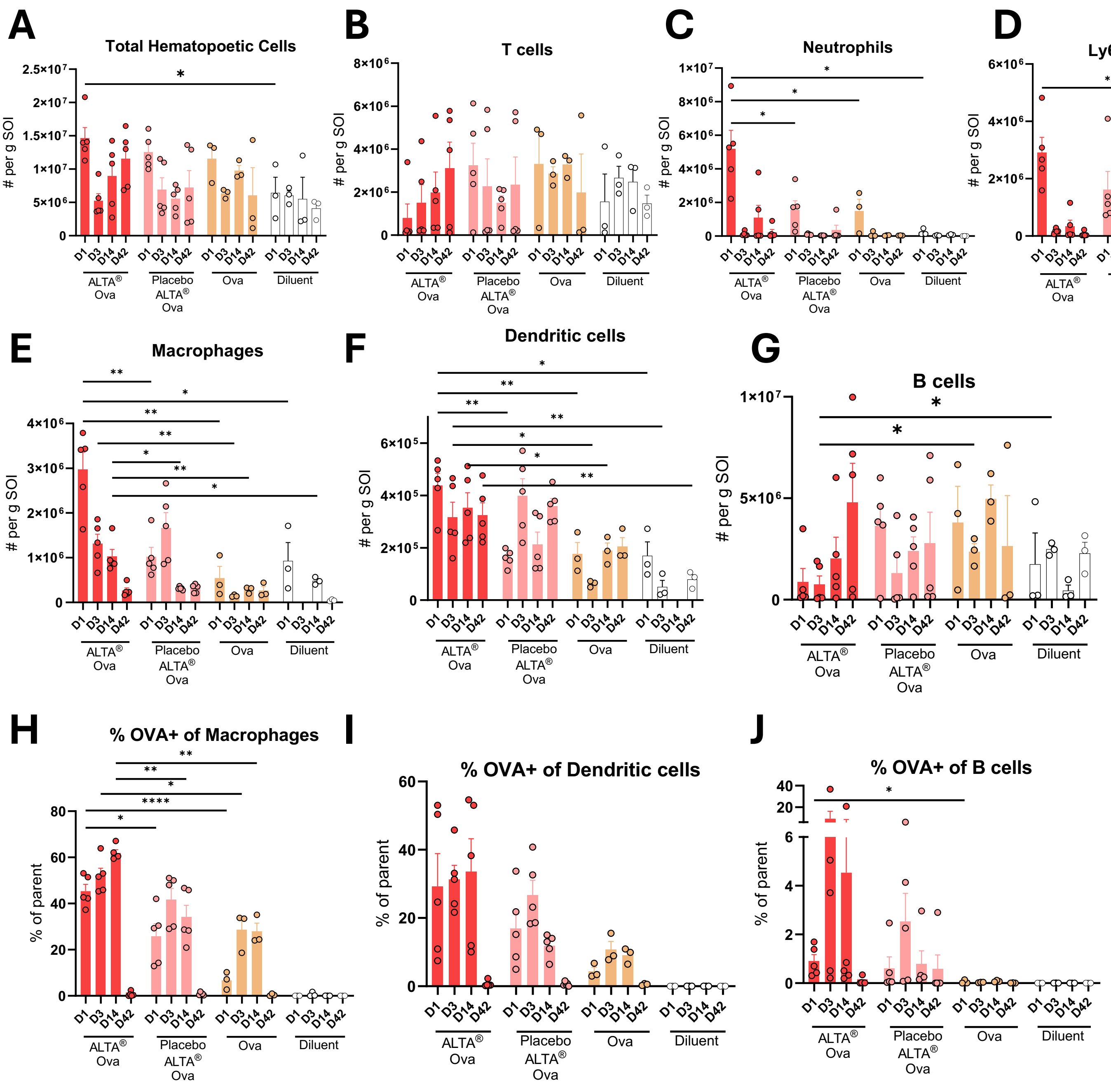

**Supplementary Figure 5. Immunization with ALTA® particles recruits immune cells to the site of injection and increases antigen capture.**

C57BL6 mice were i.m. immunized with OVA-IVISense680 dye conjugate formulated using ALTA® platform (ALTA® OVA) or OVA-IVISense680 dye conjugate mixed in diluent with placebo ALTA® particles formulated without antigen (Placebo ALTA®). Liquid control groups were injected with liquid OVA-IVISense680 or diluent. All groups except diluent control were administered 820 ng OVA dose. At days 1, 3, 7, and 42, muscle tissue containing site of injection was harvested and analyzed by flow cytometry. (**A -G**). Cell numbers per gram muscle tissue of main immune cell subsets at the SOI. Immune subsets were identified as described in supplemental gating hierarchy (Supplementary Figure 4). The percent of CD45+ was determined for each immune cell subset and cell number per gram muscle was determined as follows: [(%CD45+ of live cells / 100) * (% subset of CD45+/ 100)]/grams muscle mass. (**H** – **J**). The percentage of OVA+ events within macrophages, DCs, and B cells at the SOI 1, 3, 7, 42 days post treatments. Antigen-presenting cell populations were gated on singlets, live cells, CD45+, CD3- subsets ahead of the shown plots (**H**). Macrophages were identified as CD45+, CD11b+, Ly6G-, Ly6c int,-, and F4/80+. **(I).** DCs were identified as CD45+, I-A/I-E+, $CD11c^{hi}$ (**J**)**.** B cells were identified as CD45+, I-A/I-E+, B220+. Shown are representative data from one of two experiments. N=3-5 /group. Mean ± SEM. Unpaired t test with Welch's correction (shown are comparisons between select groups (ALTA® OVA vs other groups) at matching time points). $*p \leq 0.05$, $**p \leq 0.01$, $***p \leq 0.001$, $****p \leq 0.0001$.

### Supplementary Figure 6

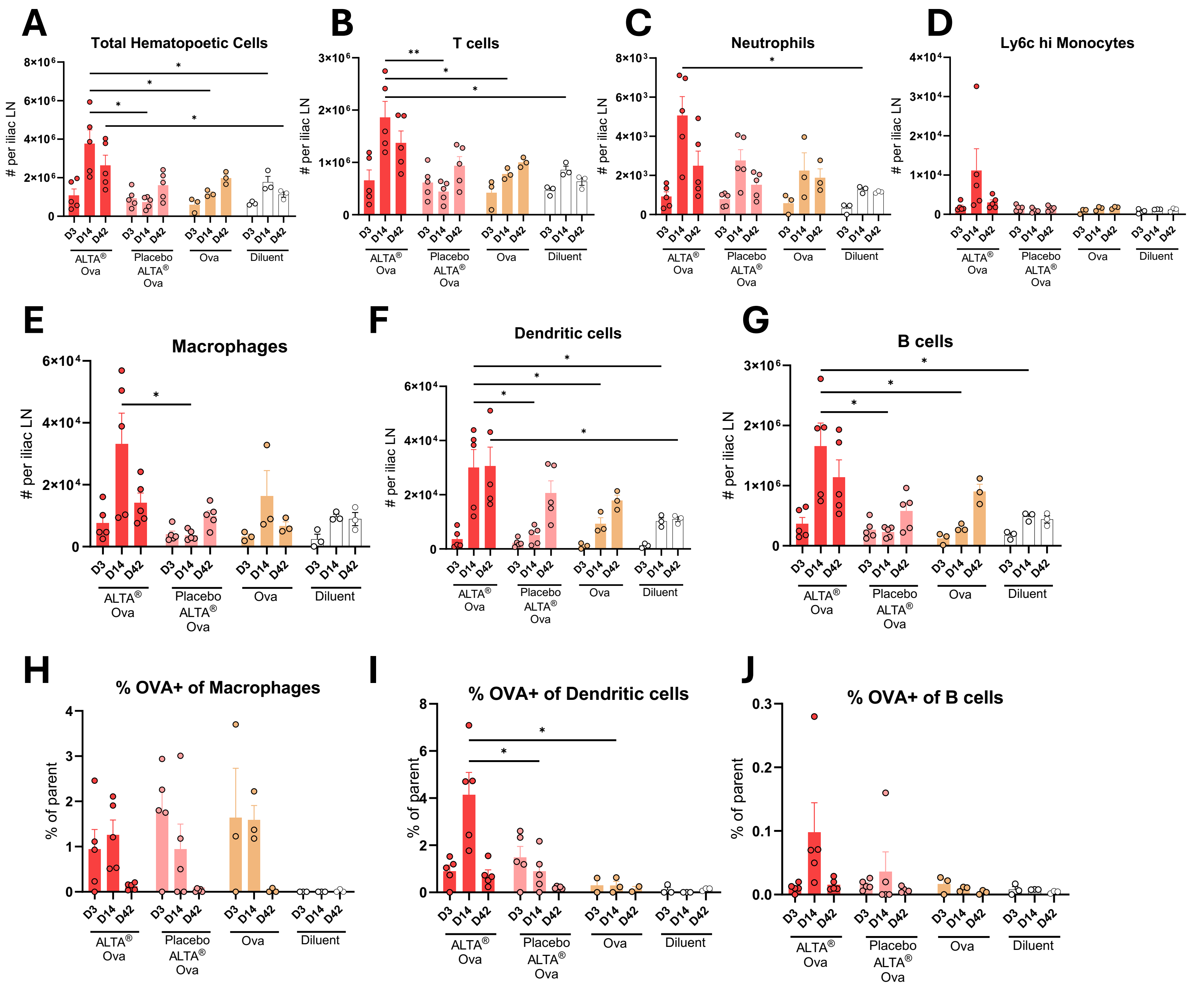

**Supplementary Figure 6. Antigen containment within ALTA® particles results in an increased number of antigen-positive cells in the dLN.**

C57BL6 mice were i.m. immunized with OVA-IVISense680 dye conjugate formulated using ALTA® platform (ALTA® OVA) or OVA-IVISense680 dye conjugate mixed in diluent with placebo ALTA® particles formulated without antigen (Placebo ALTA®). Liquid control groups were injected with liquid OVA-IVISense680 or diluent. All groups except diluent control were administered 820 ng OVA dose. At days 3,7, and 42, muscle tissue containing site of injection was harvested and analyzed by flow cytometry. (**A**-**G**) Cell numbers of main immune cell subsets per iliac LN. Immune subsets were identified as described in supplemental gating hierarchy (Supplementary Figure 4). The percent of CD45+ was determined for each subset and absolute cell counts per iliac lymph node were determined as follows: (%CD45+ of live cells / 100) * (% subset of CD45+ / 100). (**H**-**J**) The percent of OVA+ events within macrophages, DCs, B cells in iliac LN at day 3, 7, and 42 post treatments. Antigen-presenting cell populations were gated on singlets, live cells, CD45+, CD3- subsets ahead of the shown plots. (**H**) Macrophages were identified as CD45+, CD11b+, Ly6G-, Ly6c int/-, and F4/80. **(I**) DCs were identified as CD45+, I-A/I-E+, $CD11c^{hi}$. (**J)** B cells were identified as CD45+, I-A/I-E+, B220+. Shown are representative data from one of two experiments. N=3-5 /group. Mean ± SEM. Unpaired t test with Welch's correction (shown are comparisons between select groups (ALTA® OVA vs other groups) at matching time points). $*p \leq 0.05$, $**p \leq 0.01$.

#### Supplementary Figure 7

**A**

| | APIs per dose in formulation | | | | |
|---|---|---|---|---|---|
| µg per dose | **9x** | **3x** | **1x** | **0.33x** | **0.11x** |
| CPS, µg/dose | 1.84 | 0.61 | 0.21 | 0.068 | 0.023 |
| Hcp1, µg/dose | 4.5 | 1.5 | 0.5 | 0.167 | 0.056 |
| CpG ODN 2006, µg/dose | 2.25 | 0.75 | 0.25 | 0.083 | 0.028 |
| Expected Aluminum, Liquid control, µg/dose | 20.7 (from Alhyd rogel) | 6.9 (from Alhyd rogel) | 2.3 (from Alhyd rogel) | 0.77 (from Alhyd rogel) | 0.26 (from Alhyd rogel) |
| Expected Aluminum, 50-coat ALTA®, µg/dose | 11.58 | 3.86 | 1.29 | 0.43 | 0.14 |

**B**

| Group: | High Adjuvant Liquid Control | Matched Adjuvant Liquid Control | ALTA® |
|---|---|---|---|
| CPS, µg/dose | 0.38 | 0.38 | 0.38 |
| Hcp1, µg/dose | 0.76 | 0.76 | 0.76 |
| CpG ODN 2006, µg/dose | 10 | 0.38 | 0.38 |
| Aluminum, µg/dose | 86.5 (from Alhydrogel®) | 1.2 (from Alhydrogel®) | 1.2 (from ALD coating) |

**Supplementary Figure 7. Dose information for Figure 9.**

A combination of antigens derived from *B. pseudomallei (*CPS-CRM and Hcp1) and ODN 2006 were spray-dried and ALD coated to generate 50-coat ALTA® particles. (**A**) C57BL6 mice were s.c.. immunized with a range of antigen doses and CpG ODN 2006 formulated using ALTA® platform. Control mice were immunized with outlined doses of liquid antigens and adjuvants. Table describing amount (µg) of antigen and adjuvant per dose. (**B**) ALTA® platform formulated antigens and ODN 2006 and liquid controls were administered s.c. on day 0 and 28. Both liquid control groups received equivalent to ALTA® formulation antigen doses, but different doses of adjuvants (10 µg ODN 2006 + 250 µg Alhydrogel®/dose; 0.38 µg ODN 2006 + 3.5 µg Alhydrogel®/ dose).
